# Multiplex labeling and manipulation of endogenous neuronal proteins using sequential CRISPR/Cas9 gene editing

**DOI:** 10.1101/2022.01.02.474730

**Authors:** Wouter J Droogers, Jelmer Willems, Harold D MacGillavry, Arthur PH de Jong

## Abstract

Recent advances in CRISPR/Cas9-mediated knock-in methods enable labeling of individual endogenous proteins with fluorophores, to determine their spatiotemporal expression in intact biological preparations. However, multiplex knock-in methods remain limited, particularly in postmitotic cells, due to a high degree of crosstalk between genome editing events. We present Conditional Activation of Knock-in Expression (CAKE), which delivers efficient, flexible and accurate multiplex genome editing in neurons. CAKE is based on sequential gRNA expression operated by a Cre- or Flp-recombinase to control the time window for genomic integration of each donor sequence, which diminishes crosstalk between genome editing events. Importantly, CAKE is compatible with multiple CRISPR/Cas9 strategies, and we show the utilization of CAKE for co-localization of various endogenous proteins, including synaptic scaffolds, ion channels and neurotransmitter receptor subunits. Knock-in efficacy was highly sensitive to DNA vector amount, while knock-in crosstalk was dependent on the rate of donor DNA integration and timing of Cre activation. We applied CAKE to study the co-distribution of endogenous synaptic proteins using dual-color single-molecule localization microscopy, and we introduced dimerization modules to acutely control synaptic receptor dynamics in living neurons. Taken together, CAKE is a versatile method for multiplex protein labeling, enabling accurate detection, precise localization and acute manipulation of endogenous proteins in single cells.

## Introduction

The spatiotemporal distribution of proteins dictates virtually all functions of cells, and the accurate detection of endogenous proteins is an essential strategy in cell biological research (Choquet, Sainlos and Sibarita, 2021). Because protein overexpression and antibody labeling have significant limitations in accuracy and specificity, there is a pressing need to develop novel techniques that detect endogenous proteins in biological preparations. Recent CRISPR/Cas9- based genome editing methods have addressed this need by inserting (fluorescent) tags in specific genes, creating knock-ins, and now make it possible to reliably detect endogenous protein distribution with fluorescence microscopy in a wide variety of biological preparations (Auer *et al*., 2014; Nakade *et al*., 2014; Mikuni *et al*., 2016; Schmid-Burgk *et al*., 2016; Suzuki *et al*., 2016; Artegiani *et al*., 2020). However, simultaneous labeling of multiple protein species in individual cells remains challenging with commonly used genome editing methods, particularly in post-mitotic cells such as neurons. We reasoned that such genetic tools are mandatory to study the co-distribution of proteins, and would present an elegant approach to manipulate the distribution and dynamics of endogenous proteins.

The fact that neurons are postmitotic cells severely complicates both simplex and multiplex genome editing strategies: it prevents the isolation and expansion of desired clones to create isogenic cell lines and precludes multiple independent rounds of gene modification. Furthermore, insertion of the donor DNA using the highly accurate homology-directed repair (HDR) pathway predominantly occurs in the S/G2 phases of mitosis (Orthwein *et al*., 2015), and is strongly disfavored in nondividing cells. While successful genomic insertion of epitope tags using HDR in neurons has been reported (Nishiyama, Mikuni and Yasuda, 2017; Matsuda and Oinuma, 2019), most neuronal knock-in methods instead utilize the more efficient, but error-prone non-homologous end joining (NHEJ) mechanism which remains active in postmitotic cells. Most NHEJ-based methods target the coding sequence of genes (Schmid-Burgk *et al*., 2016; Suzuki *et al*., 2016; Gao *et al*., 2019; Willems *et al*., 2020), but more recent strategies replace endogenous exons (Danner *et al*., 2021; Fang *et al*., 2021) or introduce novel exons in intronic sequences (Zhong *et al*., 2021) to mitigate the effects of indel mutations. Indels can also be reduced in neurons using microhomology-mediated end joining (Yao *et al*., 2017).

Although these methods faithfully label individual proteins in neurons, multiplex epitope tagging using CRISPR/Cas9 has remained challenging. While achievable with HDR, its efficacy is generally too low for routine use (Mikuni *et al*., 2016; Matsuda and Oinuma, 2019). NHEJ in turn, operates without homology between donor DNA and target locus, and therefore the donor DNA can indiscriminately integrate in any double-stranded break (DSB), leading to a high degree of donor integration in the incorrect locus (i.e. crosstalk, Fig. 1A). Gao *et al*., 2019 circumvented this problem by creating donor DNAs that prevent protein labeling when inserted in the incorrect gene. This strategy successfully generated double knock-ins, but has restrictions on the location of the protein tag, and will generate null mutations for N-terminally labeled proteins when the incorrect donor is integrated (Gao *et al*., 2019).

**Figure. 1.**
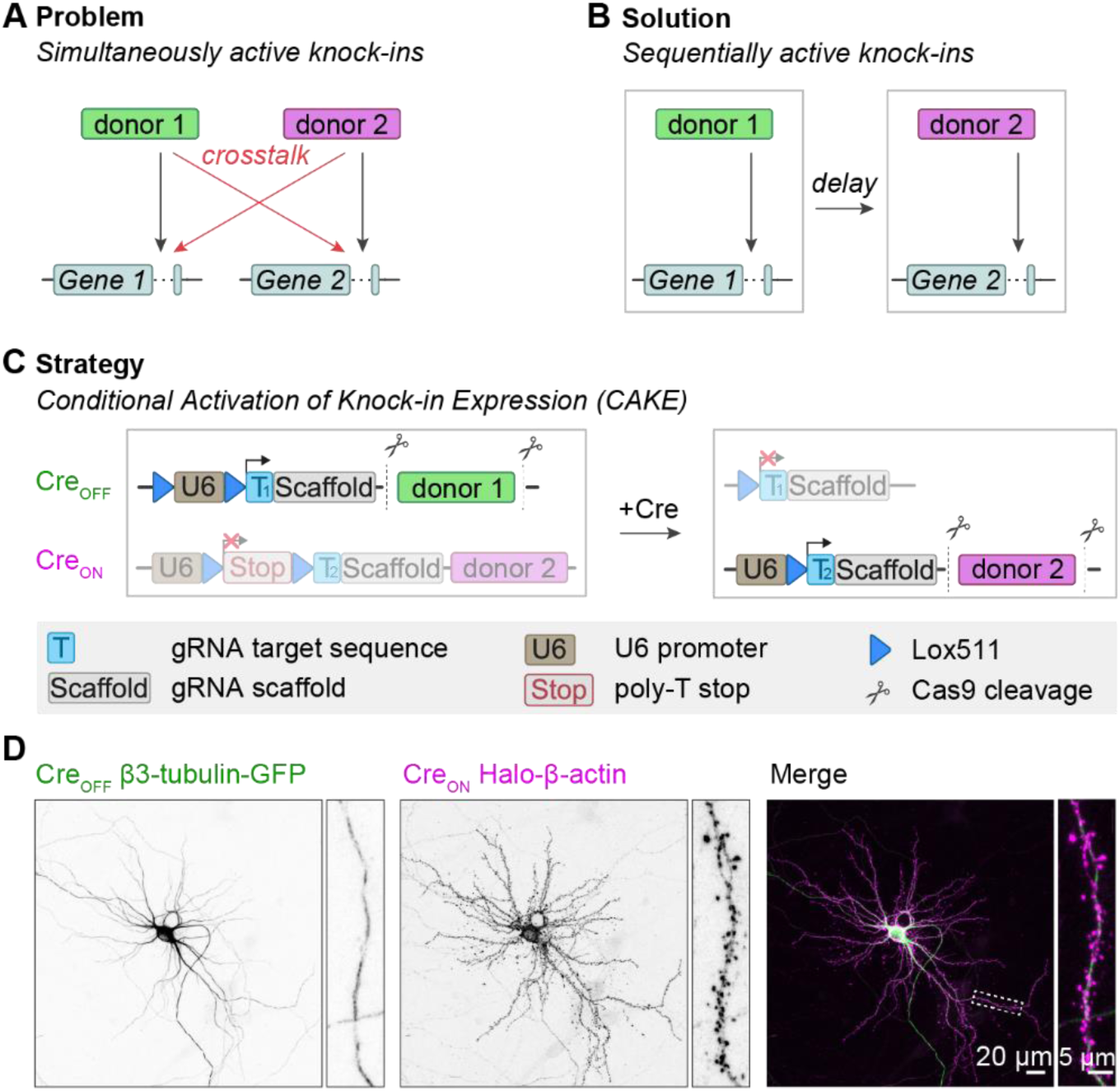
Multiplex labeling of endogenous proteins using CAKE. **A.** Illustration of the problem of multiplex knock-in strategies based on NHEJ. With simultaneous editing of multiple genes, donor DNAs can be integrated in either allele, leading to crosstalk. **B.** Proposed solution for multiplex knock-ins. By introducing a delay between two genome editing events, crosstalk can be avoided. CAKE is designed to control the delay between the two events, using Cre- or Flp-recombinase. **C.** CAKE strategy. The Cre_OFF_ vector is active in the absence of Cre, leading to editing of the first target gene, and removal of the Cre_OFF_ donor for subsequent genomic integration. Upon addition of Cre, gRNA expression from Cre_OFF_ is deactivated, and gRNA expression from Cre_ON_ vector is enabled for editing the second target gene. All donors contain a fluorophore or epitope tag flanked by a PAM- and target sequence (not shown, see Fig. S3B and Willems *et al*., 2020). **D.** Example confocal image of a Cre_OFF_ β3-tubulin-GFP and Cre_ON_ Halo-β-actin knock-in. 20 µL lenti-Cre was added at DIV 7, and cells were fixed at DIV 14.

We reasoned that crosstalk between donor DNAs could be diminished by separating genome editing events in time (Fig. 1B, also see Chylinski *et al*., 2019). Using our NHEJ-based Open Resource for the Application of Neuronal Genome Editing (ORANGE) toolbox we previously achieved this for a small number of genes with a mechanism we dubbed Conditional Activation of Knock-in Expression (CAKE) (Willems *et al*., 2020). In this approach, a GFP-2A-Cre donor sequence was fused to the first gene, which after successful knock-in switched on the expression of the second knock-in vector (Fig. S1A). However, we did observe crosstalk between the loci for some knock-in combinations, suggesting insufficient control over the delay between the two genome editing events (Willems *et al*., 2020).

To overcome this, we implemented major improvements of our CAKE strategy that result in reproducible multiplex genome editing in neuronal preparations. We demonstrate that, with Cre-dependent knock-in vectors and precise timing of Cre-recombinase activation, a high rate of correct double knock-in cells can be attained, while strongly diminishing crosstalk. Furthermore, we applied CAKE to study and manipulate the positioning of multiple endogenous proteins simultaneously in individual cells, providing a versatile method to resolve complex biological questions.

## Results

### CAKE creates double knock-ins in neurons

We reasoned that accurate multiplex knock-ins in neurons could be achieved by separating genome editing events in time using a Cre-dependent conditional activation mechanism (Fig. 1A-B). In a previous study we achieved this by fusing GFP-2A-Cre to the first gene to activate a second knock-in construct with a Cre-dependent single guide RNA (gRNA; Willems *et al*., 2020; Fig. S1A). This mechanism successfully yielded double knock-in cells for a variety of gene combinations, illustrating the potential of sequential gene editing (Willems *et al*., 2020). However, appreciable crosstalk between the knock-ins still occurred, suggesting we had insufficient control over the delay between genome editing events (Fig. S1C).

Here, we introduce two major improvements of this CAKE strategy (Fig. 1C). First, to obtain full control over the switch from the first to the second gRNA, we separately introduced Cre expression. We either used lentiviral infection of a Cre-expressing vector or lipofection of a 4OH-tamoxifen-inducible Cre-expressing construct. Second, to reduce crosstalk, we redesigned the first knock-in vector such that gRNA expression of the first vector is switched off by Cre, effectively limiting further editing of the first targeted locus (also see Chylinski *et al*., 2019). We refer to the first knock-in vector as Cre_OFF_ (gRNA expression is switched off by Cre), and the second knock-in vector as Cre_ON_ (gRNA expression switched on by Cre, see Fig. 1B). This sequential knock-in strategy yielded a mosaic of fluorescent cells, with cells positive for the Cre_OFF_ or Cre_ON_ knock-ins (Fig. S2) and a fraction of double knock-in cells that are positive for both the Cre_ON_ and Cre_OFF_ knock-ins (Fig. 1D).

We first compared the updated CAKE strategy with our previous GFP-2A-Cre based method (Willems *et al*., 2020), using knock-in constructs for β3-tubulin and β-actin, delivered to cultured rat hippocampal neurons with lipofectamine (Fig. S1). Since the distribution patterns of β3-tubulin and β-actin in neurons are well-known to be segregated in different subcellular compartments, this allowed us to easily quantify knock-in efficacy and accuracy. We systematically counted all fluorescent cells per coverslip, scoring them as a correct knock-in cell for β3-tubulin-GFP or Halo-β-actin; a double knock-in cell, or, if donor crosstalk had occurred, as an incorrect knock-in cell. Strikingly, while both methods lead to a similar number of single and double knock-in cells, the updated strategy nearly completely abolished crosstalk between knock-ins (Fig. S1D). Furthermore, we noted that for some genes GFP-2A-Cre fusion resulted in reduced expression levels, probably due to the increase in mRNA length (Willems *et al*., 2020). Importantly, no such effect was found for the improved CAKE method (Fig. S1E). Thus, CAKE faithfully created double knock-ins in cultured hippocampal neurons.

### CAKE can be applied to multiple gene combinations

To test if we could generalize the application of CAKE to other gene pairs, we generated a set of Cre_OFF_ and Cre_ON_ knock-in vectors targeting a diversity of neuronal proteins. These knock-ins include synaptic scaffolding proteins (PSD95, Homer1 and MPP2) and neurotransmitter receptor subunits (GluA1, GluN1). We also added voltage- and Ca^2+^-gated ion channels (Ca_V_2.3, SK2, BK), where the limited availability of antibodies has hampered (co-)localization analysis in neurons. Similar to β3-tubulin and β-actin knock-in vectors, these CAKE combinations yielded mosaic fluorescent labeling in cultured neurons. Importantly, we identified multiple double knock-in cells for many gene combinations (Fig. 2A). This illustrates the potential for CAKE to study the spatiotemporal co-expression of a wide range of proteins.

**Figure. 2.**
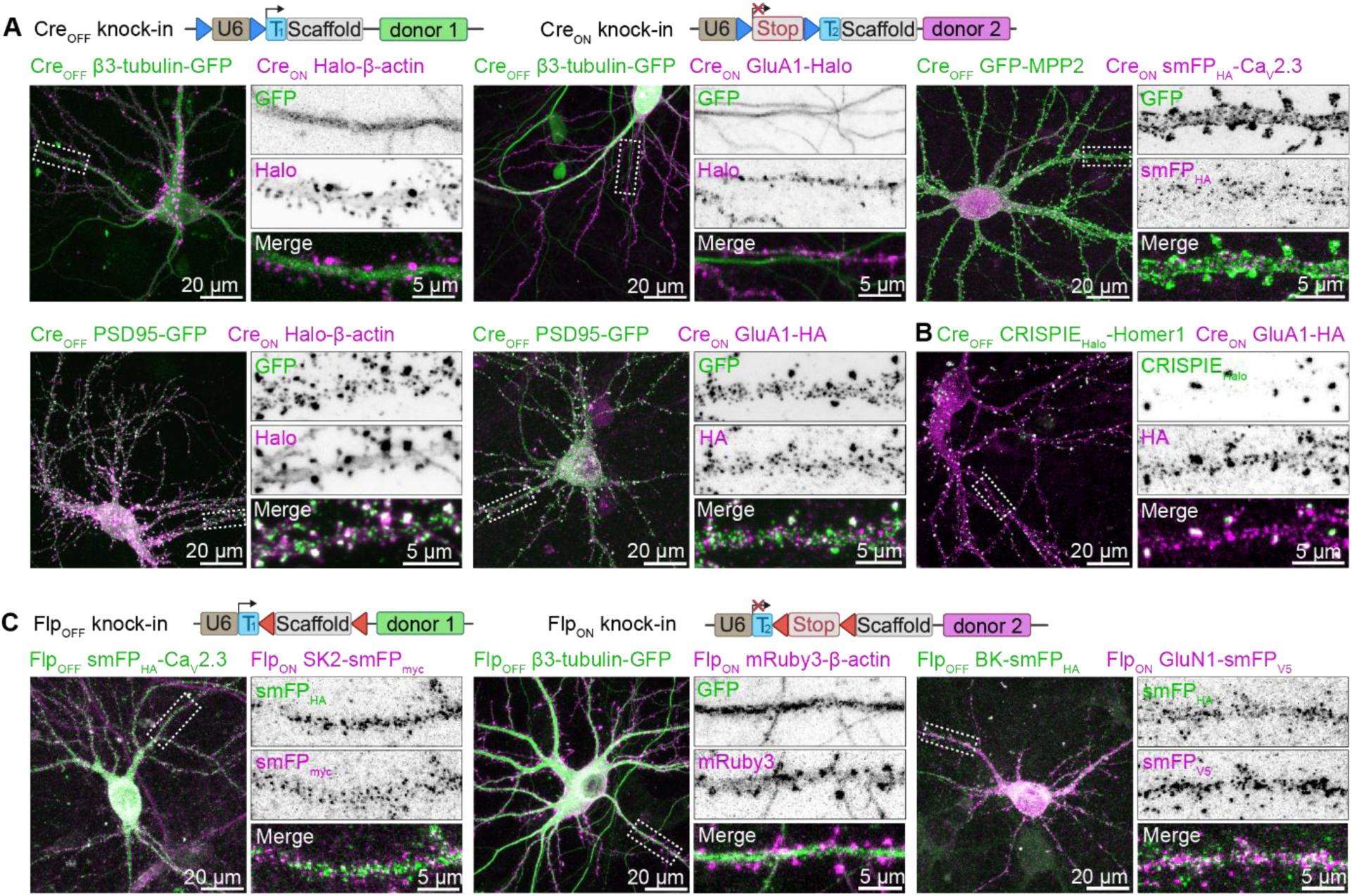
CAKE generates double knock-ins for a variety of genes. **A.** Top: Overview of Cre_OFF_ and Cre_ON_ knock-in constructs. Bottom: example confocal microscopy images of CAKE knock-in cells controlled by Cre-recombinase. For Cre_OFF_ β3- tubulin-GFP / Cre_ON_ GluA1-Halo, lenti GFP-Cre was used, which labels infected nuclei. The remaining examples were obtained with lenti-Cre without GFP, or with ER^T2^-Cre-ER^T2^. Lentivirus or 100 nM 4OH-tamoxifen was added at DIV 7. Cells were fixed at DIV 14. **B.** Example confocal image of Cre_OFF_ CRISPIE_Halo_-Homer1 and Cre_ON_ GluA1-HA double knock-in cells. CRISPIE_Halo_ donor DNA is inserted into intron 1 of Homer1 as a novel exon (see Fig. S4 and Zhong *et al*., 2021). Double knock-in was obtained using ER^T2^-Cre-ER^T2^ and 100 nM 4OH-tamoxifen was added at DIV 7. Cells were fixed at DIV 14. **C.** Top: Overview of Flp_OFF_ and Flp_ON_ knock-in constructs. Bottom: example confocal microscopy images of CAKE knock-ins controlled by lenti-Flp^O^-recombinase, added on DIV 7 Cells were fixed at DIV 14.

### CAKE is compatible with multiple CRISPR-Cas9 knock-in strategies

Because CAKE is based on sequential gRNA expression, we reasoned that our method should be compatible with other recently developed CRISPR/Cas9 knock-in strategies (e.g. Fang *et al*., 2021; Gao *et al*., 2019; Schmid-Burgk *et al*., 2016; Zhong *et al*., 2021). To assess the flexibility of CAKE, we implemented the CRISPR-mediated insertion of exon (CRISPIE) approach, which introduces designer exons in intronic sequences to mitigate the effect of indel mutations (Zhong *et al*., 2021). We inserted an exon containing Halo into the first intron of *Homer1* by flanking the donor DNA with splicing acceptor and donor sites (Fig. S4). The resulting Cre_OFF_ vector was successfully combined with a Cre_ON_ ORANGE vector to attain double knock-in cells (Fig. 2B). Thus, various NHEJ-based CRISPR-Cas9 methods can be adopted and combined with CAKE to create multiplex knock-ins.

### Controlling CAKE with Flp-recombinase

To extend the utility of CAKE, we created CAKE vectors that are controlled by Flp-recombinase (Flp_OFF_ and Flp_ON_, Fig. S3 and Fig. 2B). The Frt and stop codon sequence were contained within the gRNA, which was reported to have a higher efficacy compared to integration in the U6 promoter (Chylinski *et al*., 2019). The switch between Flp_OFF_ and Flp_ON_ gRNA expression was controlled using a lentivirus expressing Flp^O^ (lenti-Flp). Flp-controlled CAKE performed comparable to Cre-controlled CAKE, and resulted in single and double knock-in cells for a variety of gene combinations (Fig. 2C). Finally, we developed a Cre_ON_ Flp_OFF_ knock-in vector, enabling intersectional activation of gRNA expression (Fig. S3A). Thus, CAKE can be performed with both Cre and Flp-recombinase.

### Knock-in efficacy is modulated by donor DNA levels

Initially, the number of double knock-in cells per coverslip was too low for many applications (Fig. S1D). Therefore, we next set out a series of experiments to increase the number of double knock-in cells per sample, using Cre_OFF_ β3-tubulin-GFP and Cre_ON_ Halo-β-actin knock-in vectors. In early experiments we noticed that vector amounts used in transfection affected knock-in efficacy. To thoroughly test this, we systematically varied the amount of the Cre_OFF_ knock-in vector in our transfection mixture between 20 and 197 fmol (which equals to 50 to 500 ng DNA) per coverslip, while keeping Cre_ON_ fixed at 178 fmol, and scored all fluorescent cells per coverslip at day in vitro (DIV) 14. Strikingly, we found a strong inverse relationship between Cre_OFF_ β3-tubulin-GFP vector amount and the number of β3-tubulin-GFP positive cells (Fig. 3A, *p* = 0.006, 2-way ANOVA). Furthermore, lower Cre_OFF_ amounts also increased the number of Halo-β-actin positive cells, even though we kept the amount of Cre_ON_ vector constant in all conditions (Fig. 3A, *p* = 0.04. 2-way ANOVA). This interplay suggests competition between the two knock-ins, which continues after the Cre-dependent switch has occurred. Together with an increase in single knock-in cells, we observed a strong increase in double knock-in cells to between 5 and 8 cells per coverslip at the lowest Cre_OFF_ β3-tubulin-GFP amount (Fig. 3A, *p* = 0.001, 2-way ANOVA). Crucially, the number of incorrect knock-in neurons (i.e. donor crosstalk for one of the targeted genes) remained low (1-2 cells per coverslip, <1% of all knock-in cells), although the absolute number slightly increased with lower DNA amount (*p* = 0.02, 2-way ANOVA). In the same experiment, we tested if the timing of Cre infection (infection at DIV 3, 7 or 9) affects the number of knock-in neurons. In contrast to the strong effect of vector amount, the timing of Cre expression did not influence the number of single or double knock-in neurons (Fig. 3A, Cre_OFF_ *p* = 0.90, Cre_ON_ *p* = 0.88, double knock-in *p* = 0.49, incorrect knock-in *p* = 0.50, 2-way ANOVA), suggesting that either knock-in efficacy for this gene combination is insensitive to Cre timing, or that the onset of lentiviral-mediated Cre expression is too slow to observe an effect. Similar results were obtained for Cre_OFF_ GluA1-GFP and Cre_ON_ PSD95-Halo, though the sensitivity for DNA amount appeared to differ between individual knock-in constructs (Fig. S5A). We then tested if further reduction of vector amount enhanced knock-in efficacy, and varied the amount of both the Cre_OFF_ and Cre_ON_ vectors. Here, we found that the optimum for Cre_OFF_ β3-tubulin-GFP is around 3.9 fmol DNA per coverslip. Cre_ON_ Halo-β-actin performed best around 36 fmol per coverslip, and efficacy dropped steeply below 3.6 fmol (Fig. 3B). This is in line with our previous observation that increased amounts of Cre_OFF_ knock-in vector negatively affect the Cre_ON_ knock-in efficacy (Fig. 3A).

**Figure 3.**
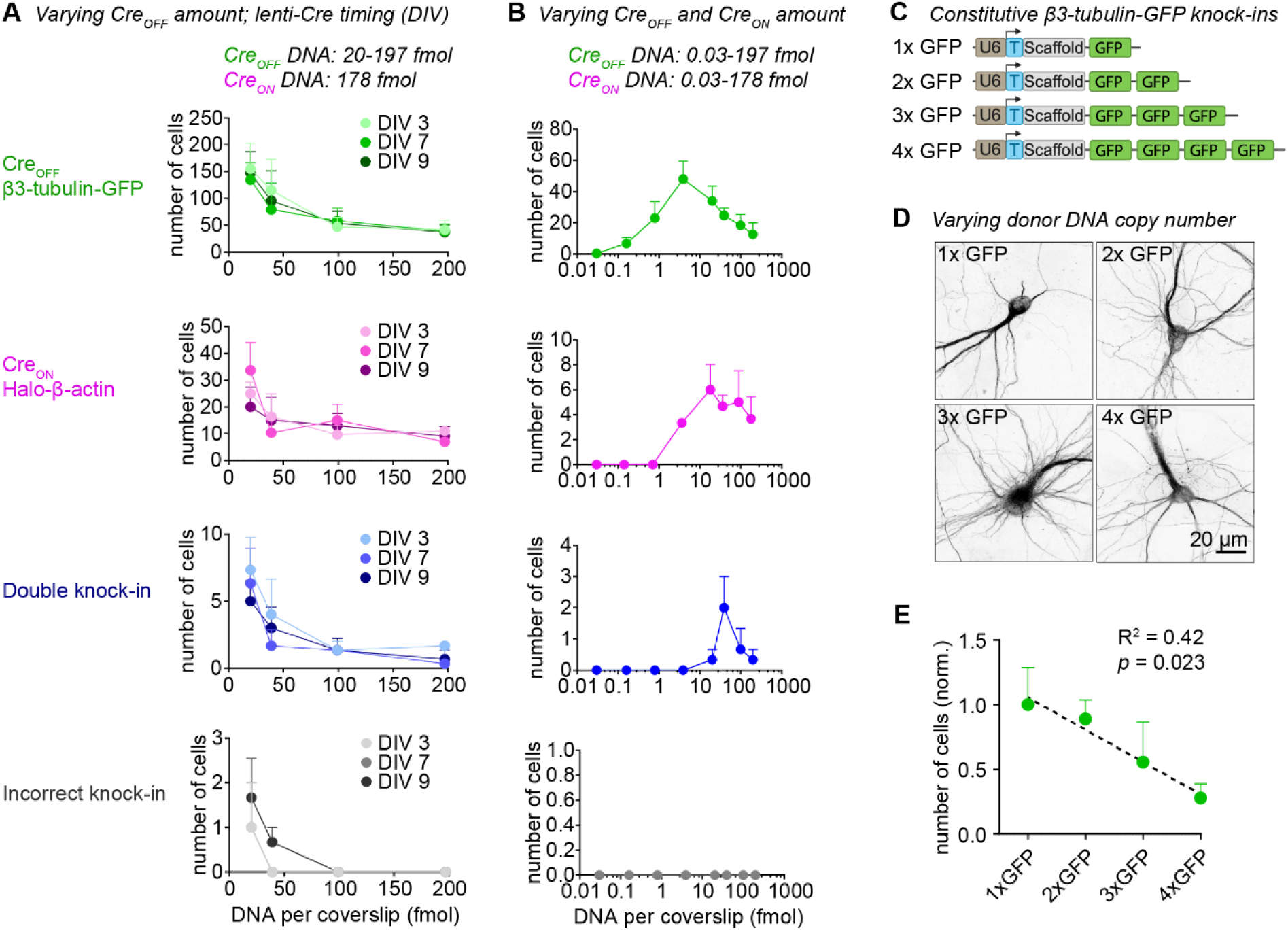
Donor DNA amount controls knock-in efficacy. **A.** Number of fluorescent cells per coverslip for each knock-in, as a function of Cre_OFF_ β3-Tubulin-GFP vector amount. 20 µL lenti-Cre was added at DIV 3, 7 or 9. n = 3 coverslips, N = 3 independent cultures. **B.** Number of fluorescent cells per coverslip for each knock-in. Both the amount of Cre_OFF_ and Cre_ON_ vector were varied; 20 µL lenti-Cre was added at DIV 7. n = 3 coverslips, N = 3 independent cultures. **C.** pORANGE β3-tubulin knock-in constructs used to titrate the amount of donor DNA. Each GFP donor has its own PAM and target sequence (not shown), and thus every GFP donor can be cleaved independently from the vector. **D.** Example confocal images of β3-tubulin-GFP knock-in cells using 1 to 4 GFP donors per vector. **E.** Number of fluorescent cells as a function of number of GFP donors. Data was normalized to average number of knock-ins in the 1xGFP condition. R^2^ = 0.42, *p* = 0.023, model linear regression (dotted black line). n = 3 coverslips, N = 3 independent cultures.

We hypothesized that the inverse relationship between DNA amount and knock-in efficacy is due to competition between donor DNAs. To test if increasing donor DNA copies per cell decreases knock-in efficacy, we generated ORANGE β3-tubulin knock-in vectors with 1, 2, 3 or 4 independent copies of the donor DNA (Fig. 3C-E). Using these vectors, we found a clear negative correlation between the number of donor DNA copies and the number of β3-tubulin-GFP positive cells per coverslip (Fig. 3E, *p* = 0.023, R^2^ = 0.42).

Finally, to separate effects of gRNA and donor DNA levels, we replaced the Cre_OFF_ β3-tubulin-GFP vector with a Cre_OFF_ gRNA vector and a minicircle GFP donor (Fig. S5B). Previous studies found that minicircle donors often outperform large donor plasmids, likely because Cas9-mediated cleavage of the donor plasmid can lead to integration of the vector backbone (Schmid-Burgk *et al*., 2016; Suzuki *et al*., 2016; Danner *et al*., 2021). We found that, at high donor levels, minicircle donor DNA performed similar to ORANGE donor plasmids, but knock-in efficacy was reduced at lower minicircle levels (Fig. S5C). The reason for this reduction is unclear, but we cannot exclude that at low amounts of minicircle DNA the transfection efficacy is reduced. Similar to knock-in vector-delivered donor, the high amounts of the (Cre_OFF_-activated) minicircle donor also decreased the efficacy of the Cre_ON_ Halo-β-actin knock-in (Fig. S5C). Importantly, increasing Cre_OFF_ gRNA expression level had no effect on knock-in efficacy for both the Cre_OFF_ and Cre_ON_ knock-in. Thus, under our experimental conditions, a single vector containing both the gRNA and donor DNA leads to the highest knock-in efficacy. Taken together, we conclude that donor DNA levels modulate knock-in efficacy.

### Tamoxifen-inducible Cre controls Cre_ON_ knock-in efficacy

In the experiments described above, we consistently observed a lower efficacy of Cre_ON_ compared to the Cre_OFF_ knock-ins. To improve the efficacy of Cre_ON_ knock-ins, we switched to 4OH-tamoxifen inducible Cre (ER^T2^-Cre-ER^T2^), which ensures rapid onset of Cre activation (Matsuda and Cepko, 2007) and superior control over timing, compared to lentiviral-mediated Cre expression. As found by others (e.g. Forni *et al*., 2006; Higashi *et al*., 2009), we observed that strong, sustained activation of ER^T2^-Cre-ER^T2^ appears toxic to neurons, which resulted in fewer single and double knock-in cells (Fig. S6B). To prevent toxicity, we reduced the vector encoding for ER^T2^-Cre-ER^T2^ to 2 fmol per coverslip, and we developed a self-inactivating Cre, by flanking ER^T2^-Cre-ER^T2^ with LoxP sites (ER^T2^-Cre-ER^T2^ lox, adapted from Pfeifer *et al*., 2001; Silver and Livingston, 2001). Under these conditions, 4OH-tamoxifen induced a dose-dependent increase in Cre_ON_ and double knock-ins (Cre_ON_ *p* = 0.0001; double knock-in *p* = 0.0001, 2-way ANOVA), without affecting the number of Cre_OFF_ knock-in cells (Fig. 4B, *p* = 0.70, 2-way ANOVA). A ten-fold increase in ER^T2^-Cre-ER^T2^ lox vector (to 20 fmol) increased the number of Cre_ON_ at low 4OH-tamoxifen concentrations (Fig. 4B, *p* = 0.025, 2-way ANOVA post-hoc comparison), but did not further increase efficacy at 100 or 1000 nM 4-OH-tamoxifen. Overall, no statistical difference was observed between ER^T2^-Cre-ER^T2^ and ER^T2^-Cre-ER_T2_ lox conditions (Cre_OFF_ knock-ins *p* = 0.44, Cre_ON_ knock-ins *p* = 0.10, double knock-ins *p* = 0.98, 2-way ANOVA). The number of incorrect knock-ins remained low (about 1 per coverslip at 1000 nM 4OH-tamoxifen), and was also similar between the Cre conditions (Fig. 4B, *p* = 0.80, 2-way ANOVA). Thus, at low vector concentrations ER^T2^-Cre-ER^T2^ accurately controls gRNA expression, while minimizing cytotoxicity.

**Figure 4.**
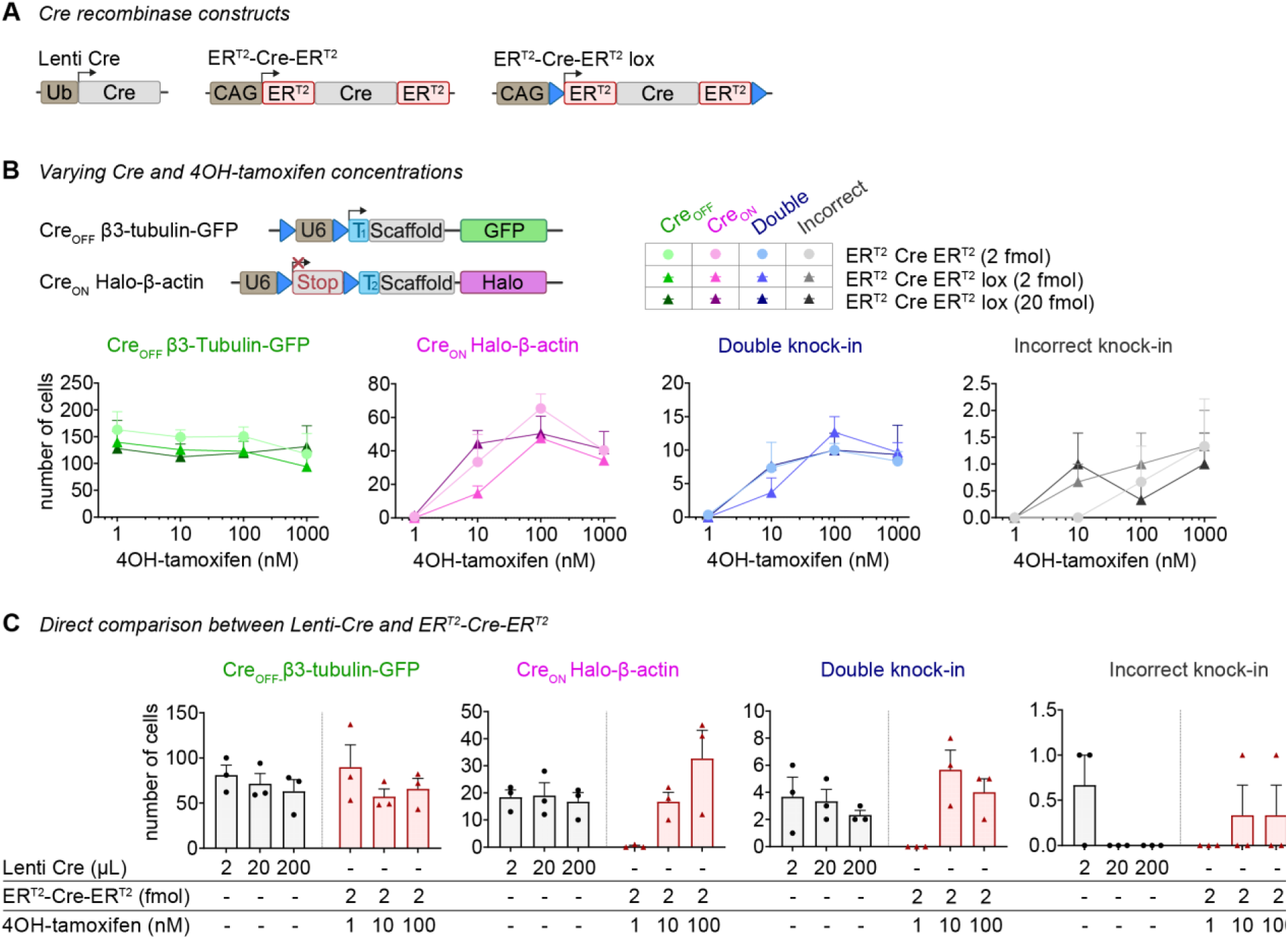
Comparison of methods to deliver and activate Cre-recombinase. **A.** Overview of Cre-recombinase constructs. **B.** Number of fluorescent knock-in cells per coverslip using ER^T2^-Cre-ER^T2^ constructs. 4OH-tamoxifen was added at the indicated concentration at DIV 7. n = 3 coverslips, N = 3 independent cultures. **C.** Comparison of Lenti-Cre and ER^T2^-Cre-ER^T2^. Lenti-Cre and 4OH-tamoxifen were added at DIV 7. n = 3 coverslips, N = 3 independent cultures.

Finally, we directly compared lenti-Cre and ER^T2^-Cre-ER^T2^, and found that ER^T2^-Cre-ER^T2^ did not boost the number of double knock-in neurons significantly (Fig. 4C). Taken together, multiple methods for Cre delivery and activation can be used to control CAKE, without obvious differences in the number of knock-in neurons. All subsequent experiments are based on ER^T2^-Cre-ER^T2^, activated on DIV 7 with 100 nM 4OH-tamoxifen.

### Crosstalk between knock-ins is dependent on timing of Cre activation

While testing several CAKE combinations, we noticed that one particular combination, using Cre_OFF_ GluA1-Halo and Cre_ON_ Flp_OFF_ smFP_V5_-Ca_V_2.3, showed an unusually high rate of donor crosstalk (Fig. 5A-C). Specifically, ∼90% of these incorrect knock-ins were smFP_V5_-positive in a staining pattern expected for GluA1 (ON-to-OFF crosstalk). To understand why the CAKE mechanism failed to prevent crosstalk in this experiment, we decided to investigate this further.

**Figure 5.**
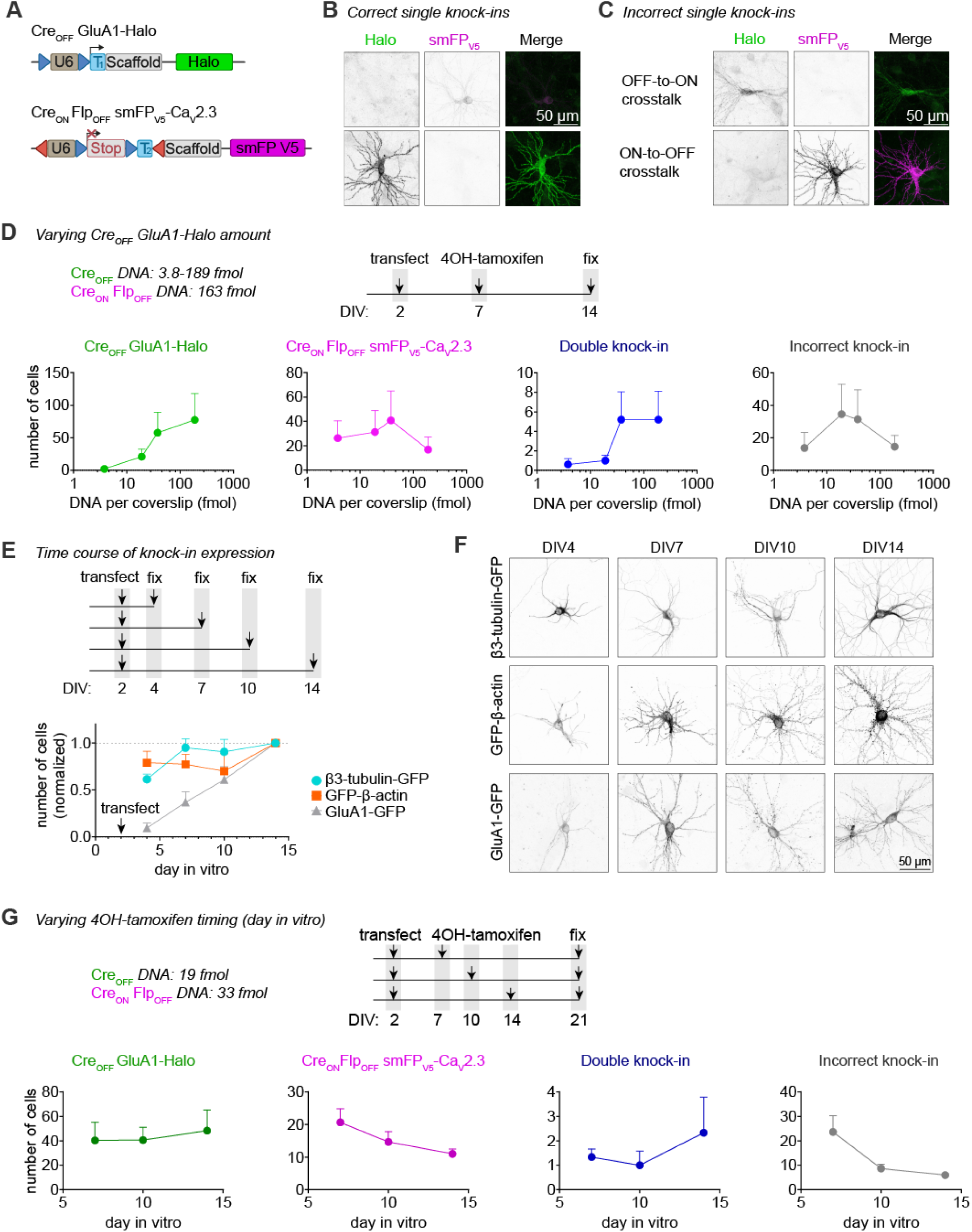
Delaying Cre activation reduces knock-in crosstalk. **A.** Overview of used DNA constructs. **B-C.** Example images of correct (B) and incorrect (C) knock-in neurons. Cells were incubated with 100 nM 4OH-tamoxifen at DIV 7, and fixed at DIV 14. Images in B and C were taken with same acquisition settings to illustrate differences in expression level between Ca_V_2.3 (top) and GluA1 (bottom). **D.** Top: experimental design. Bottom: Number of fluorescent cells per coverslip for each knock-in, as a function of Cre_OFF_ GluA1-Halo vector amount. 100 nM 4OH-tamoxifen was added at DIV 7. Cells were fixed at DIV 14. n = 5 coverslips, N = 5 independent cultures. **E.** Top: experimental design. Experiment was performed without Cre or 4OH-tamoxifen. Bottom: Number of fluorescent cells per coverslip at different timepoints, normalized to the number of cells at DIV 14. n = 4 coverslips, N = 4 independent cultures. **F.** Example confocal microscopy images of cells fixed at different DIVs from E. Image acquisition settings were kept identical per knock-in. **G.** Top: experimental design. Bottom: Number of fluorescent cells per coverslip for each knock-in, as a function of the DIV at which 100 nM 4OH-tamoxifen was added. n = 3 coverslips, N = 3 independent cultures.

Several features of these two knock-in constructs favor the detection of ON-to-OFF crosstalk. Firstly, both gRNAs target the same position in the reading frame of their respective target genes (frame +1). Secondly, Cre_OFF_ GluA1-Halo targets the GluA1 C-terminus with a stop codon in the Halo donor, while Cre_ON_ Flp_OFF_ smFP_V5_-Ca_V_2.3 targets the Ca_V_2.3 N-terminus.

Thus, ON-to-OFF crosstalk would lead to an in-frame addition of smFP_V5_ in *Gria1*, while OFF-to-ON crosstalk would introduce an early stop codon in *Cacna1e*, likely preventing protein expression from the allele.

To test if crosstalk could be prevented, we first titrated the amount of Cre_OFF_ GluA1-Halo vector. Unexpectedly, we found that higher Cre_OFF_ GluA1-Halo load increased the number of GluA1-Halo knock-in cells (even though the effect was not obvious in statistical analysis due to high variability between cultures (Fig. 5D, *p* = 0.21, 1-way ANOVA), the opposite of what we previously found for Cre_OFF_ β3-tubulin (Fig. 3A). Higher Cre_OFF_ GluA1-Halo load also appeared to increase the number of double knock-in cells and incorrect knock-ins (Cre_ON_ Flp_OFF_ smFP_V5_-Ca_V_2.3 *p* = 0.80, double knock-ins *p* = 0.26, incorrect knock-ins *p =* 0.63). At 38 fmol Cre_OFF_ vector, about 25% of all fluorescent cells were incorrect (predominantly ON-to-OFF crosstalk).

We hypothesized that this crosstalk is due to a low editing rate of *Gria1* compared to genes with lower rates of crosstalk (e.g. *Tubb3* and *Actb* Fig. 3), which continues after Cre activation at DIV 7. Indeed, several studies demonstrated that both the appearance of DSBs, as well as (DNA-repair dependent) indels are highly dependent on the sequence of the target locus, and that repair of DSBs may continue for multiple days (Rose *et al*., 2017; Liu *et al*., 2020; Park *et al*., 2021). To compare editing rates over time, we transfected Cre_OFF_ GFP knock-ins for β3-tubulin, β-actin and GluA1 at DIV 2, and fixed and counted the number of GFP positive cells at DIV 4, 7, 10 and 14 (Fig. 5E). For all knock-ins, we found that the number of GFP-positive cells increased over time (*p* = 1.0 x 10^-5^, 2-way ANOVA), and this effect differed between knock-in constructs (*p* = 5.0 x 10^-5^, interaction *p* = 0.012, 2-way ANOVA). Importantly, while the number of β3-tubulin and β-actin knock-in neurons increased at a similar rate (*p* = 0.81, 2-way ANOVA post-hoc comparison), the number of GluA1 knock-in neurons increased much slower (β3-tubulin vs. GluA1 *p* = 0.0006, β-actin vs GluA1 *p* = 3.5 x 10^-3^, 2-way ANOVA post-hoc comparison) and continued to increase after DIV 10.

Finally, we tested if delaying Cre activation, by delaying the addition of 4OH-tamoxifen, would reduce ON-to-OFF crosstalk (Fig. 5G). We found that delaying 4OH-tamoxifen addition had no effect on the number of single or double knock-in neurons (Cre_OFF_ GluA1-Halo *p* = 0.91, Cre_ON_ Flp_OFF_ smFP_V5_-Ca_V_2.3 *p* = 0.17, double knock-ins *p* = 0.59, 1-way ANOVA), but clearly diminished the number of incorrect knock-in cells (*p* = 0.041, 1-way ANOVA). Thus, while editing of *Gria1* is much slower compared to other genes, crosstalk can be largely reduced by delayed activation of Cre.

### CAKE enables dual-color single-molecule localization microscopy of endogenous synaptic proteins

Mapping the localization of endogenous synaptic proteins is crucial for our understanding of the brain (Choquet, Sainlos and Sibarita, 2021). In particular, deciphering which proteins regulate AMPA receptor nanoscale clustering in glutamatergic synapses will be pivotal in understanding synaptic transmission. Interestingly, AMPA receptors have been shown to be concentrated in subsynaptic PSD95 nanodomains (MacGillavry *et al*., 2013; Nair *et al*., 2013). Here, we used CAKE to uncover the nanoscale co-organization of endogenously tagged PSD95 and the AMPA receptor subunit GluA1 at glutamatergic synapses with dual-color single-molecule localization microscopy (SMLM).

As expected, endogenously tagged PSD95 and GluA1 co-localized at synapses (Fig. 6A). Using custom local density-based cluster analysis (based on Chen *et al*., 2020 and MacGillavry *et al*., 2013), we identified nanodomains of both GluA1 and PSD95 within individual synapses (Fig. 6B). Synapses contained on average more PSD95 nanodomains, but larger GluA1 nanodomains (Fig. S7A-B, nanodomain number *p* = 2.0 x 10^-4^, nanodomain diameter *p* = 6.4 x 10^-8^, unpaired *t*-test). These GluA1 nanodomains occurred on average only 14 nm closer to the center of the postsynaptic density (PSD) (Fig. 6C, *p* = 5.2 x 10^-8^, unpaired *t*-test), and the distance between PSD95 and GluA1 nanodomains only slightly increased with larger PSDs (Fig. 6D, *p* = 0.0002, R^2^ = 0.021, model linear regression). This suggests that PSD95 and GluA1 have similar synaptic topologies across synapse sizes. To further assess the co-organization of PSD95 and GluA1, we used a local density-based co-localization index (Fig. 6E and Fig. S7C; see Materials and Methods for details). Individual synapses had highly variable degrees of co-localization of PSD95 and GluA1 (Fig. S7C). In addition, the co-localization of PSD95 and GluA1 was correlated with PSD size (Fig. 6F, *p* = 1.0 x 10^-15^, R^2^ = 0.19), suggesting that stronger synapses have a tighter association between GluA1 and PSD95. Lastly, we observed that co-localization between GluA1 and PSD95 was higher inside nanodomains compared to the rest of the PSD (Fig. 6G, PSD95 *p* = 6.1 x 10^-6^, GluA1 *p* = 1.6 x 10^-7^, one-sample *t*-test), confirming previous observations (MacGillavry *et al*., 2013; Nair *et al*., 2013). In summary, CAKE allowed us to reveal the tight nanoscale co-organization of endogenous PSD95 and AMPA receptors, which correlated with synapse size.

**Figure 6.**
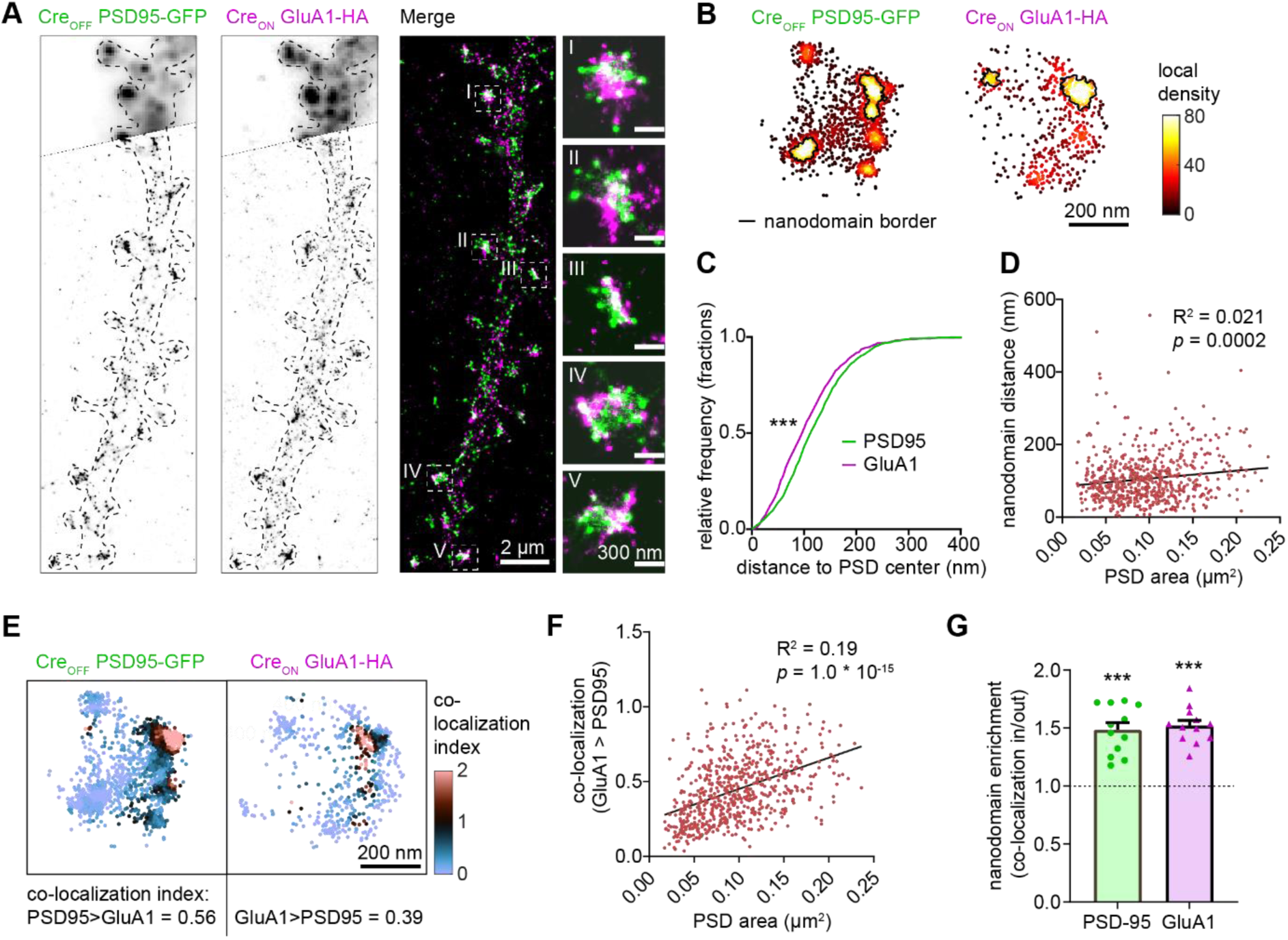
Dual-color SMLM reveals nanoscale co-organization of endogenous PSD95 and GluA1. **A.** SMLM reconstruction of PSD95-GFP and GluA1-HA. Top part of the two channels shows widefield image of the same dendrite. Pixel size of reconstruction: 12 x 12 nm. **B.** Example synapse showing the local density values of PSD95 and GluA1 localizations. Nanodomains are outlined with a black line. **C.** Nanodomain topology of PSD95 and GluA1 nanodomains. Cumulative distributions of the distance between nanodomain and PSD center for both PSD95 (green) and GluA1 (magenta). GluA1 nanodomains reside on average 14 nm closer to the PSD center. *** *p* = 5.2 * 10^-8^, Mann-Whitney test. 1285 nanodomains were analyzed for PSD95 and 774 nanodomains were analyzed for GluA1. n = 12 neurons, N = 4 independent cultures. **D.** Distance between PSD95 and GluA1 nanodomains correlates weakly with PSD size. PSD size is based on the cluster of PSD95 localizations inside the synapse (see Materials & Methods). R^2^ = 0.021, *p* = 0.0002, model linear regression (black line). 656 synapses were analyzed. n = 12 neurons, N = 4 independent cultures. **E.** Example synapse showing the co-localization index of PSD95 and GluA1 localizations. The average co-localization index of this synapse is indicated below the graph. **F.** The co-localization of GluA1 with PSD95 localizations correlates with PSD size. R^2^ = 0.192, *p* = 1.0 x 10^-15^, model linear regression (black line). 656 synapses were analyzed. n = 12 neurons, N = 5 independent cultures. **G.** PSD95 and GluA1 are both enriched in nanodomains of the other channel. Relative co-localization of PSD95 and GluA1 inside versus outside nanodomains is plotted. Dotted line represents null hypothesis (no nanodomain enrichment). PSD95 *** *p* = 6.1 x 10^-6^, GluA1 *** *p* = 1.6 x 10^-7^, one sample *t*-test.

### Acute immobilization of endogenous synaptic AMPA receptors using inducible hetero-dimerization

Labeling of two endogenous proteins simultaneously in single neurons is a powerful means to measure and modulate their (functional) co-localization, but experiments are often hampered by overexpression artifacts. For example, the amount of PSD95 as well as AMPA receptors determine synaptic strength, and overexpression of these synaptic components can result in altered synaptic function (El-Husseini *et al*., 2000; Schnell *et al*., 2002). Interestingly, precise control of lateral diffusion and synaptic exchange of AMPA receptors is important for both basal transmission and synaptic plasticity (Groc and Choquet, 2020). Thus, studying the trafficking and anchoring of AMPA receptors requires methods that accurately modulate the localization of endogenous proteins in living neurons. Here, we used CAKE to label AMPA receptors and PSD95 with inducible dimerization modules to acutely manipulate AMPA receptor anchoring at the synapse. More specifically, we generated C-terminal CAKE knock-in constructs to label GluA1 and PSD95 with the rapalog-inducible dimerization domains FRB and FKBP respectively (Fig. 7A; Kapitein *et al*., 2010). Addition of rapalog would then anchor AMPA receptors to PSD95 proteins at the synapse, thus reducing receptor exchange between the PSD and the extrasynaptic membrane (Fig. 7B).

**Figure 7.**
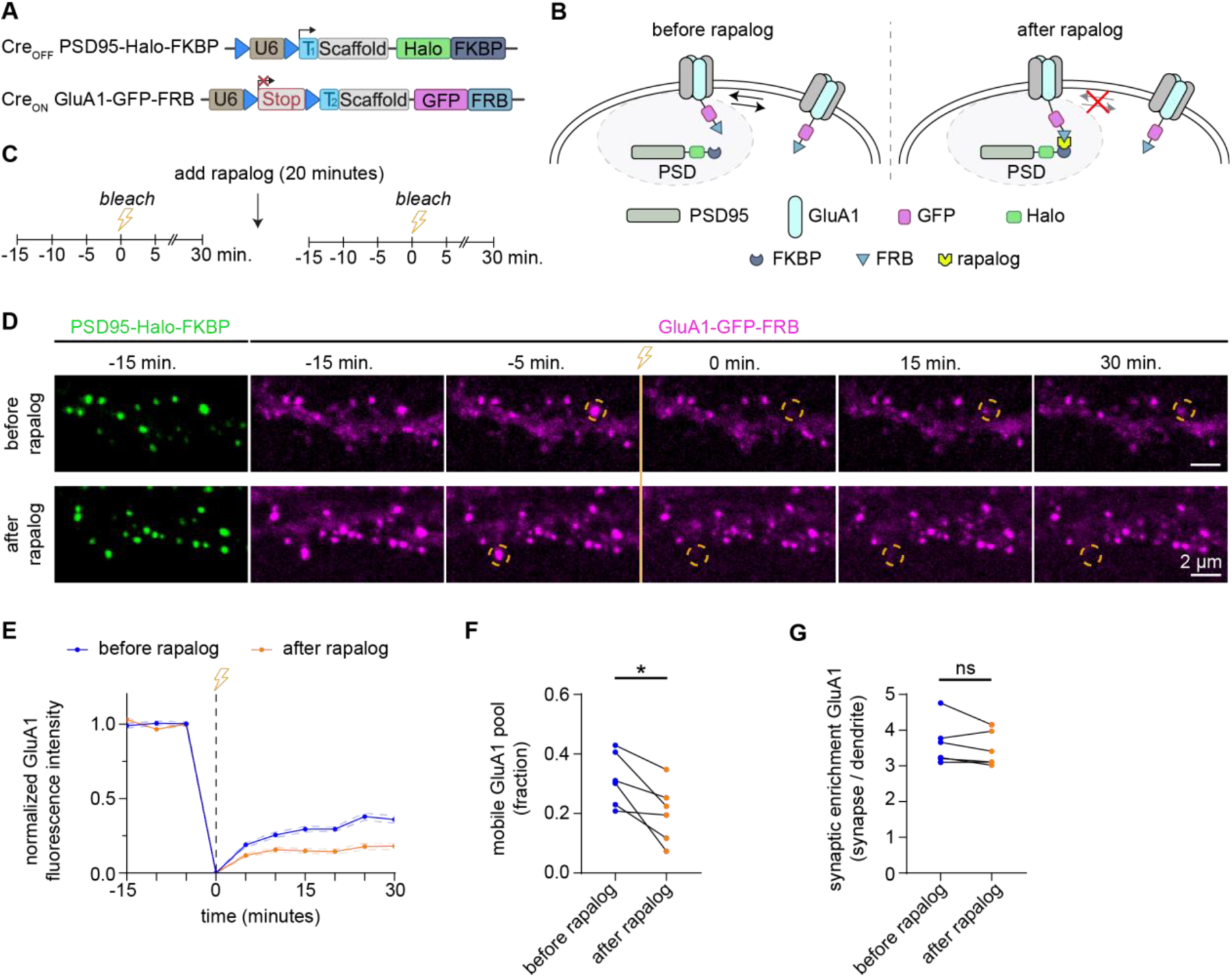
Live-cell modulation of endogenous AMPA receptor anchoring using CAKE. **A.** Overview of DNA constructs used. **B.** Graphical overview of synaptic anchoring of AMPA receptors using rapalog. FRB (fused to GluA1), binds to FKBP (fused to PSD95), preventing exchange of synaptic receptors. **C.** Imaging protocol. Neurons are imaged every 5 minutes using spinning disk confocal microscopy. FRAP is performed twice, before and after the incubation with rapalog for 20 minutes. **D.** Example FRAP acquisition before and after incubation with rapalog. GluA1-GFP-FRB in magenta and PSD95-Halo-FKBP in green. Spines indicated with orange circle are bleached just before timepoint 0 minutes. **E.** FRAP curves of spines bleached before and after incubation with rapalog. Data is normalized to the average intensity before bleaching. 63 spines before rapalog and 66 spines after rapalog were bleached. n = 6 cells, N = 5 independent cultures. **F.** Average recovery of fluorescence per neuron, averaged over the last 4 frames, reflecting the mobile pool of receptors. After rapalog, the mobile pool of receptors is less than before rapalog. * *p* = 0.018, paired t-test. n = 6 cells, N = 5 independent cultures. **G.** Synaptic enrichment of GluA1-GFP-FRB at synapses before and after rapalog incubation are similar. Synaptic enrichment is the relative fluorescence intensity at the synapse compared to the dendrite. *p* = 0.21, paired *t*-test. n = 6 cells, N = 5 independent cultures.

We performed live-cell spinning disk imaging experiments to measure the fluorescence recovery after photobleaching (FRAP) at synapses to quantify GluA1 turnover under basal conditions and after the addition of rapalog (Fig. 7C-D). Under basal conditions, the fluorescence recovery of GluA1 receptors was 0.31 ± 0.04 (Fig. 7E-F), which is consistent with previous studies (Chen *et al*., 2021; Fang *et al*., 2021). We incubated neurons with rapalog for 20 minutes, and measured FRAP dynamics on a different dendrite of the same neuron. Rapalog induced a strong decrease in fluorescence recovery (0.20 ± 0.04) of GluA1, indicating successful anchoring of AMPA receptors to PSD95 (Fig. 7E-F, *p* = 0.018, paired *t*-test). Importantly, we did not observe a decrease of GluA1 turnover in neurons that were not treated with rapalog (Fig. S8, *p* = 0.86, paired *t*-test). We found no significant change in the enrichment of GluA1 at the synapse after rapalog (Fig. 7G, *p* = 0.21, paired *t*-test). Together, these results show that CAKE allows for labeling and rapidly inducible dimerization of endogenous proteins in living neurons.

## Discussion

Accurate detection and manipulation of endogenous proteins is essential to understand cell biological processes, which motivated laboratories across cell biology to develop highly efficient CRISPR genome editing methods for endogenous epitope tagging (Auer *et al*., 2014; Nakade *et al*., 2014; Lackner *et al*., 2015; Schmid-Burgk *et al*., 2016; Suzuki *et al*., 2016; Nishiyama, Mikuni and Yasuda, 2017; Artegiani *et al*., 2020; Danner *et al*., 2021). Multiplex editing using NHEJ-based CRISPR/Cas9 methods remains limited due to the high degree of crosstalk that occurs between two knock-in loci (Gao *et al*., 2019; Willems *et al*., 2020). In the current study we present CAKE, a mechanism to diminish crosstalk between NHEJ-based CRISPR/Cas9 knock-ins using sequential activation of gRNA expression. We demonstrate that this mechanism strongly reduces crosstalk between knock-in loci, and results in dual knock-ins for a wide variety of genes. Finally, we show that CAKE facilitates super-resolution microscopy and live manipulation of multiple endogenous proteins in neuronal cells.

### Comparison of CAKE with other CRISPR/Cas9 strategies

Previous studies for NHEJ-based multiplex knock-ins had restrictions on donor DNA design and the target loci that could be combined (e.g. C-terminal or C-terminal knock-ins), and these methods could inadvertently reduce protein expression from the targeted allele (Gao *et al*., 2019; Willems *et al*., 2020). The CAKE knock-in strategy presented here lifts these limitations, and can be used with any locus and donor DNA design. Importantly, because CAKE only relies on sequential gRNA expression it is expected to be compatible with any combination of NHEJ-based knock-in modalities, including HITI (Suzuki *et al*., 2016), HiUGE (Gao *et al*., 2019), TKIT (Fang *et al*., 2021), REPLACE (Danner *et al*., 2021) and CRISPIE (Fig. 2B; Zhong et al., 2021).

### Factors that influence knock-in efficacy

The CAKE mechanism presented here creates a mosaic of Cre_ON_ and Cre_OFF_ knock-ins, and the number of double knock-in cells depends on the efficacy of each knock-in vector. Therefore, to obtain a high number of double knock-in cells, the efficacy of both the Cre_ON_ and Cre_OFF_ knock-in vector must be optimized. We identified three parameters that regulate the efficacy for single- and double knock-ins in neurons. Firstly, the efficacy of gRNAs varies widely, and even gRNAs that target sequences a few base pairs apart in the same locus can have dramatically different knock-in rates (Willems *et al*., 2020; Danner *et al*., 2021; Fang *et al*., 2021; Zhong *et al*., 2021). Thus, the efficacy of each individual gRNA must be optimized in order to increase the chance of successful multiplex labeling in neurons. gRNA performance is dependent on many factors, including the rate of DNA cleavage and repair (Rose *et al*., 2017; Liu *et al*., 2020; Park *et al*., 2021) and the propensity of the target locus for indel mutations (Rose *et al*., 2017; Shen *et al*., 2018; Liu *et al*., 2020). While some of these parameters can be predicted computationally (Doench *et al*., 2014; Shen *et al*., 2018; Park *et al*., 2021), the efficacy of each individual gRNA should be verified experimentally. We performed most experiments with gRNAs that we and others previously found to yield a high number of single knock-ins (Suzuki *et al*., 2016; Willems *et al*., 2020).

Secondly, we found knock-in efficacy to be highly sensitive to knock-in vector amount. For multiple knock-ins, we found that reducing the amount of knock-in vector increased the number of knock-in positive cells, with the optimum for β3-tubulin as little as 3.9 fmol vector DNA per coverslip. We propose that this inverse relationship is due to competition between donor DNA molecules for integration. In line with these observations, knock-in vectors with multiple donors reduced the number of β3-tubulin-GFP positive cells. Donor competition could also explain why we consistently needed more knock-in vector for the Cre_ON_ knock-in, as remaining donor from the Cre_OFF_ vector could compete for integration in the target locus after Cre activation. Importantly, for reasons incompletely understood, the optimum vector amount differs considerably between knock-in constructs. A striking example in this respect is GluA1, which requires a 10-to-50-fold high vector load to reach the maximum number of knock-in cells. We also observed that onset of GluA1 knock-in expression is much slower compared to other genes, and thus there may be a relationship between the editing rate of the targeted locus, and the amount of donor DNA required for successful integration.

Thirdly, the timing of Cre expression and activation may influence the number of incorrect knock-in cells. For Cre_OFF_ GluA1 / Cre_ON_ Ca_V_2.3 we found that delaying the activation of ER^T2^-Cre-ER^T2^ by 7 days diminished crosstalk between the knock-ins. This is in line with our observation that GluA1 knock-ins are completed at a much slower rate than other genes tested here. We did not observe an effect of infection day for lenti-Cre with Cre_OFF_ β3-tubulin and Cre_ON_ β-actin. However, the slow onset of expression from lentiviral vectors (in the order of days; Hioki *et al*., 2007) makes lentivirus a weak method to observe an effect Cre timing, in particular compared to the rapid activation of ER^T2^-Cre-ER^T2^ (Matsuda and Cepko, 2007). Additionally, donor integration in *Tubb3* appears be relatively fast, which limits the window for crosstalk to occur.

### Application of CAKE for labeling and manipulation of endogenous proteins

CAKE opens possibilities to study localization, mobility and function of endogenous proteins at multiple levels. For instance, the nanoscale organization of synaptic proteins profoundly influences information transfer at synapses (MacGillavry *et al*., 2013; Nair *et al*., 2013; Tang *et al*., 2016; Rebola *et al*., 2019), and CAKE may present an invaluable tool to decipher synapse organization using endogenous protein labeling under stringent conditions required for super resolution microscopy. Indeed, we were able to accurately determine the subsynaptic co-localization of GluA1 and PSD95, which can be used to assess the functionality and requirements of their nanoscale co-organization. To study protein-protein interactions CAKE could also be applied in combination with Förster resonance energy transfer reporters, bimolecular fluorescence complementation (Tebo and Gautier, 2019), or proximity biotinylation assays (De Munter *et al*., 2017).

Our results also illustrate the potential of CAKE to manipulate the localization, mobility and functionality of endogenous proteins, for instance by recruiting or anchoring proteins and organelles in living cells using optogenetic or chemical dimerization modules. Trafficking and sub-cellular positioning of proteins is crucial for all cellular processes, and this unique combination of techniques allows manipulation of endogenous protein dynamics in cells. For example, directed positioning of receptors (Sinnen *et al*., 2017) or even entire organelles (Van Bergeijk *et al*., 2015) have previously been shown to influence synaptic strength and neuronal development respectively.

The conditional activation of CRISPR/Cas9 knock-ins also opens new avenues for detailed analysis of endogenous proteins in individual cell types. By restricting Cre or Flp expression using cell-type specific promoters (Taniguchi *et al*., 2011; He *et al*., 2016), combined with Cre_ON_ or Flp_ON_, one could map out the spatiotemporal expression of proteins in a wide variety of cells at unprecedented precision.

Taken together, we created and validated a series of CRISPR tools for sequential genome editing to create multiplex knock-ins with NHEJ, and demonstrate the value of these tools for determining and manipulating protein distribution in neurons.

## Materials and Methods

### Ethics statement

All experiments were approved by the Dutch Animal Experiments Committee (Dier Experimenten Commissie (DEC; AVD1080020173404), performed in line with institutional guidelines of Utrecht University, and conducted in agreement with Dutch law (Wet op de Dierproeven, 1996) and European regulations (Directive 2010/63/EU). Timed pregnant Wistar rats were obtained from Janvier Labs.

### Molecular cloning

Constructs were made using standard laboratory techniques. All CAKE knock-in backbones are numbered as pORANGE CAKE (pOCx); see Fig. S3A and Table 1 for a complete overview. For most pOCx vectors, variants with and without CAG SpCas9 were created. All primer sequences are found in Table 3.

**Table 1.**
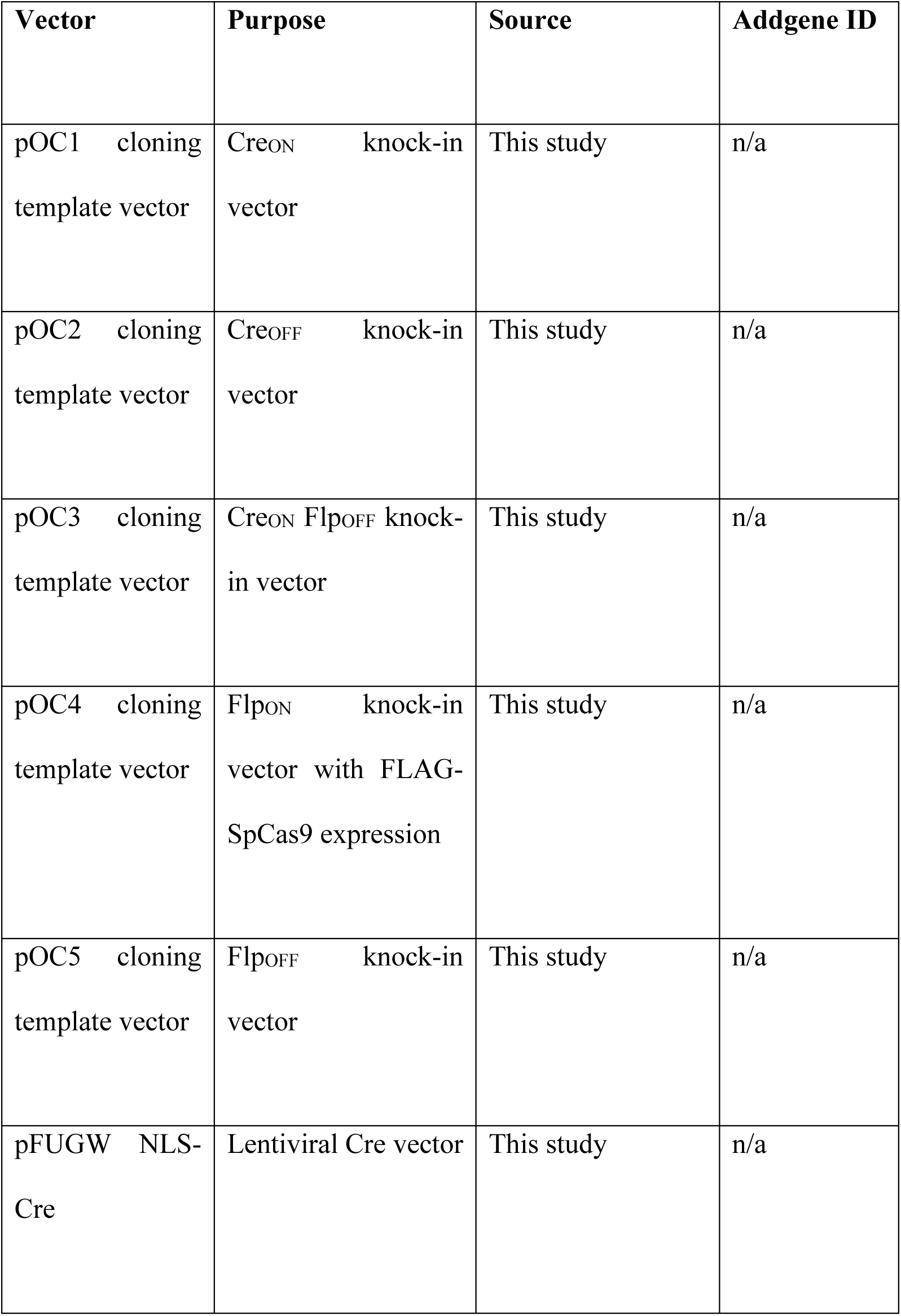
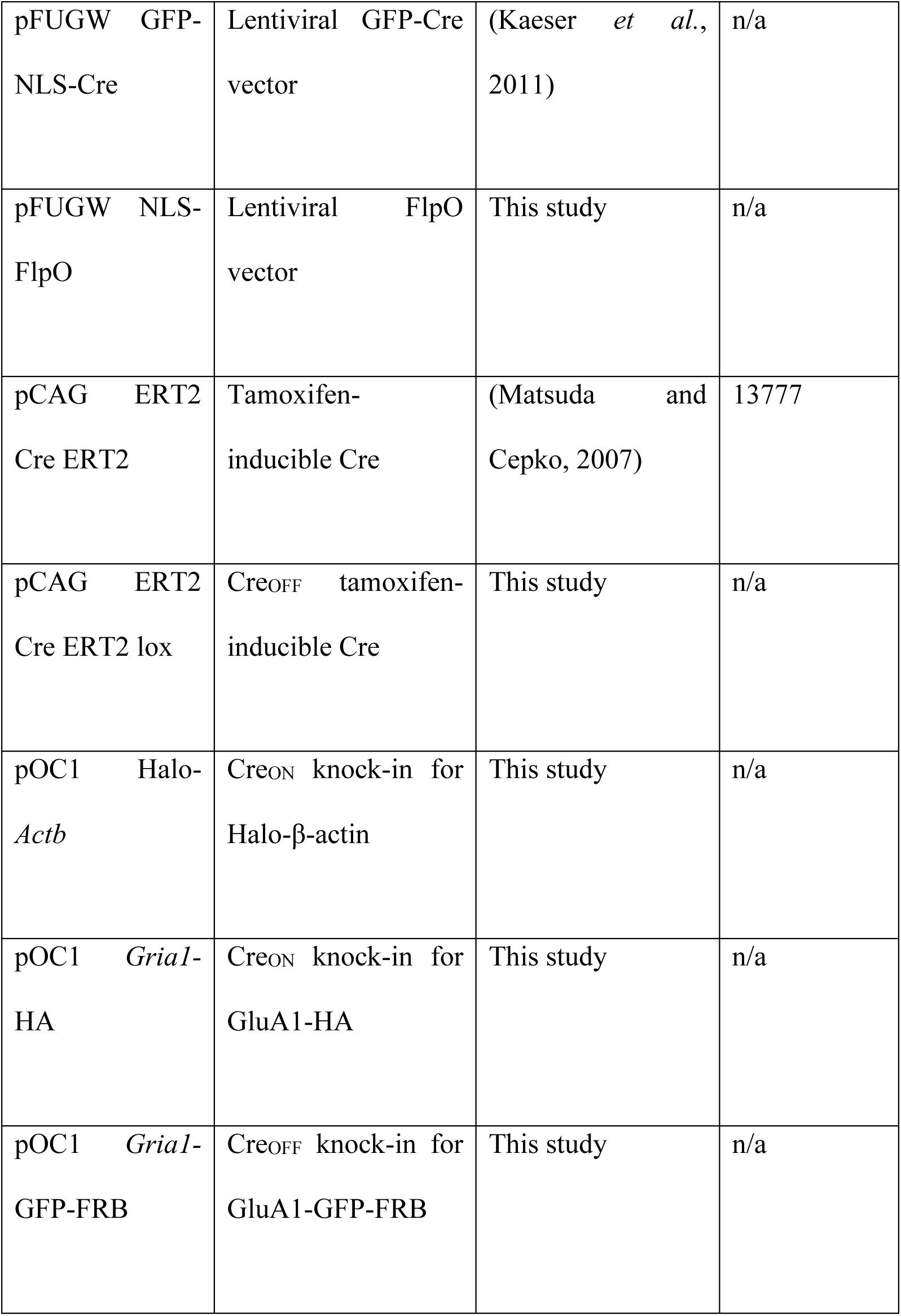
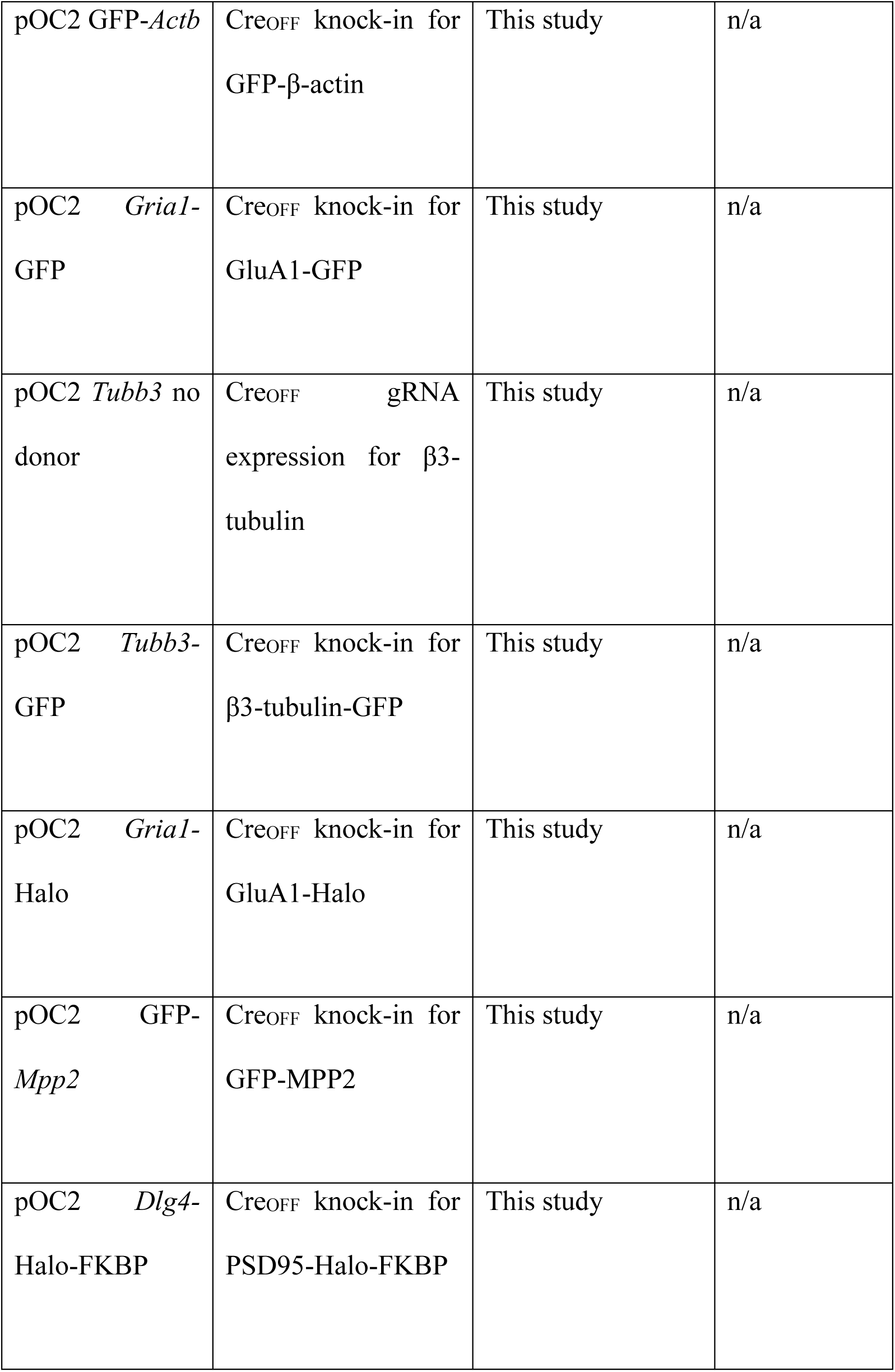
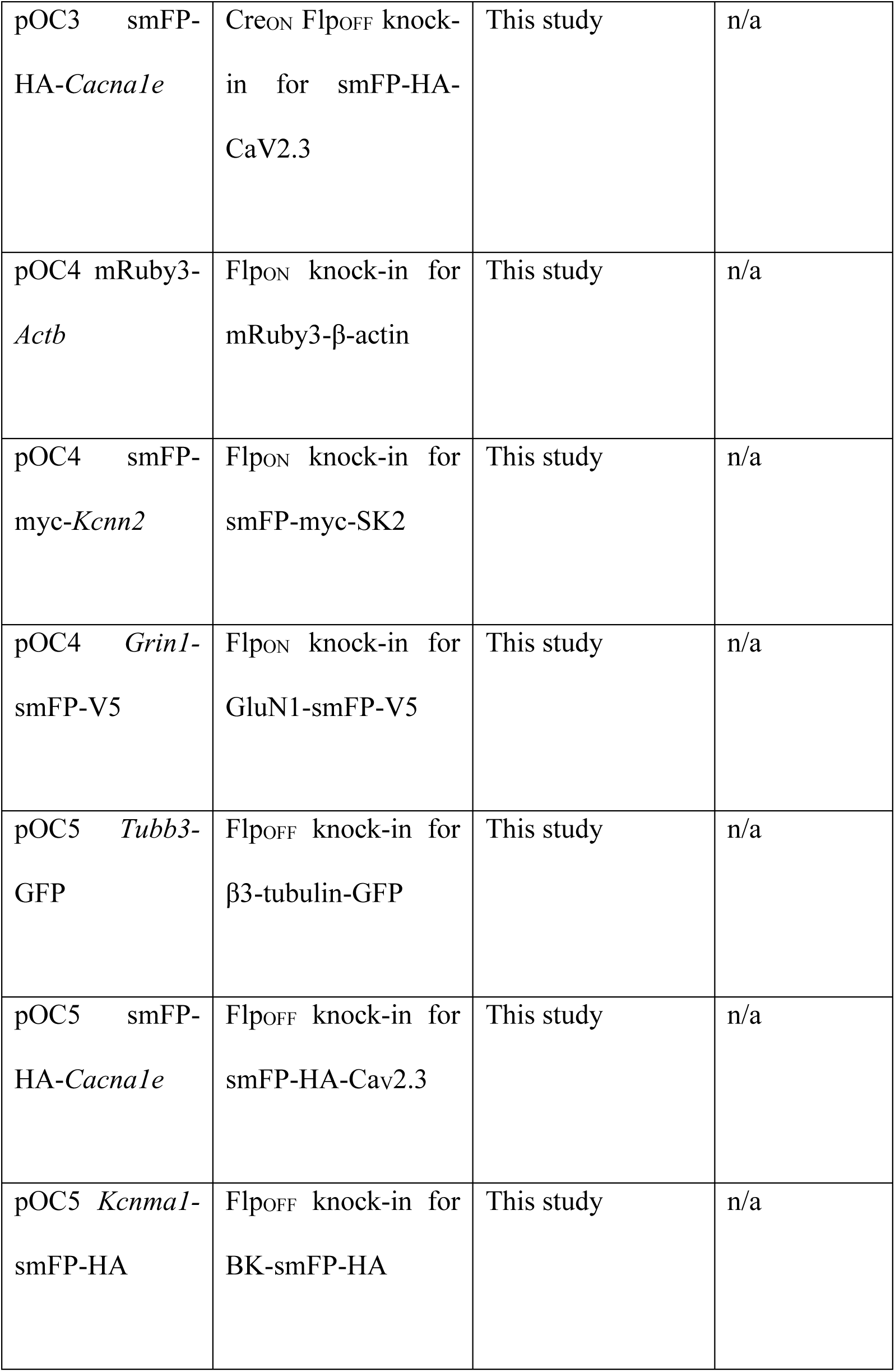
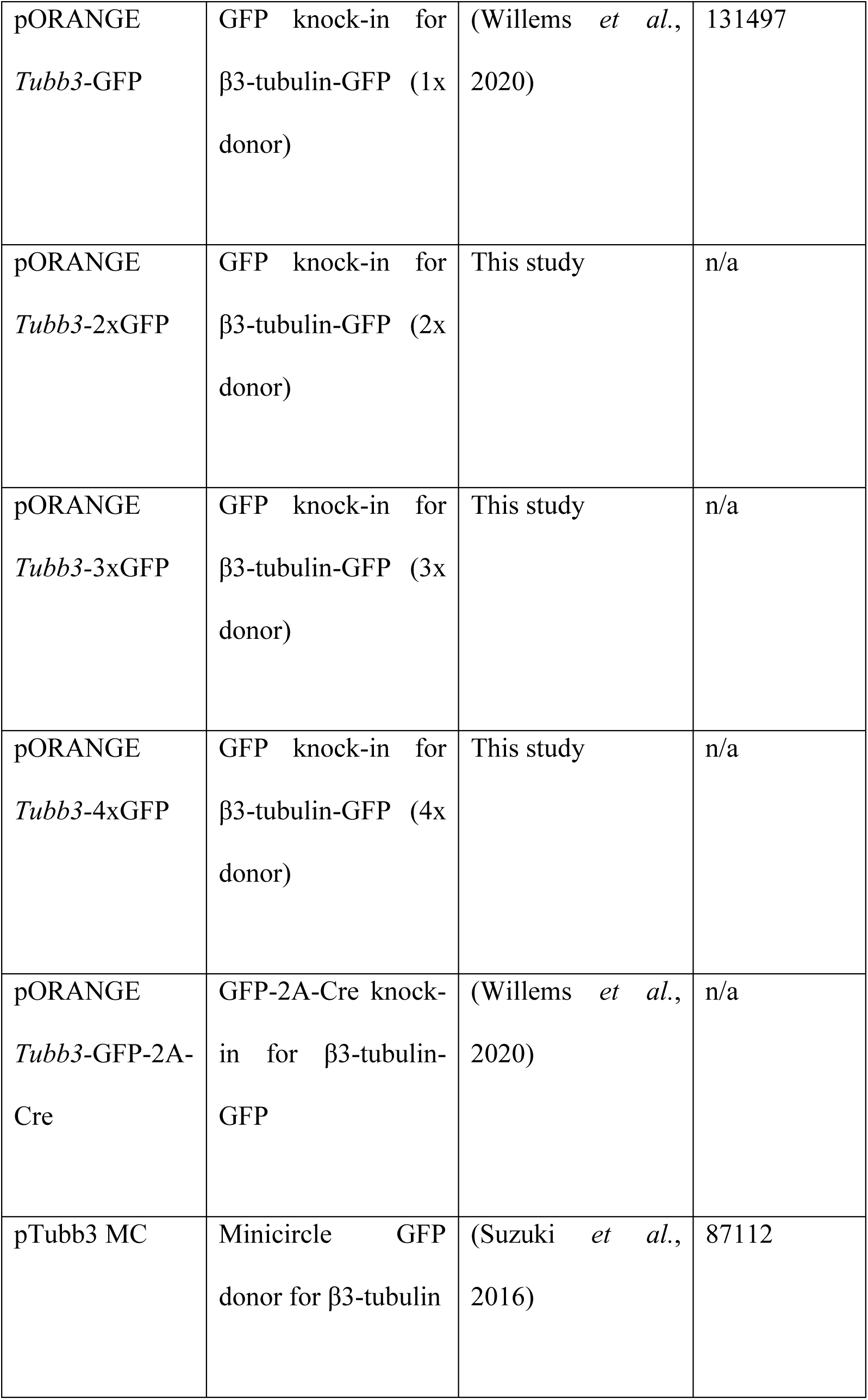
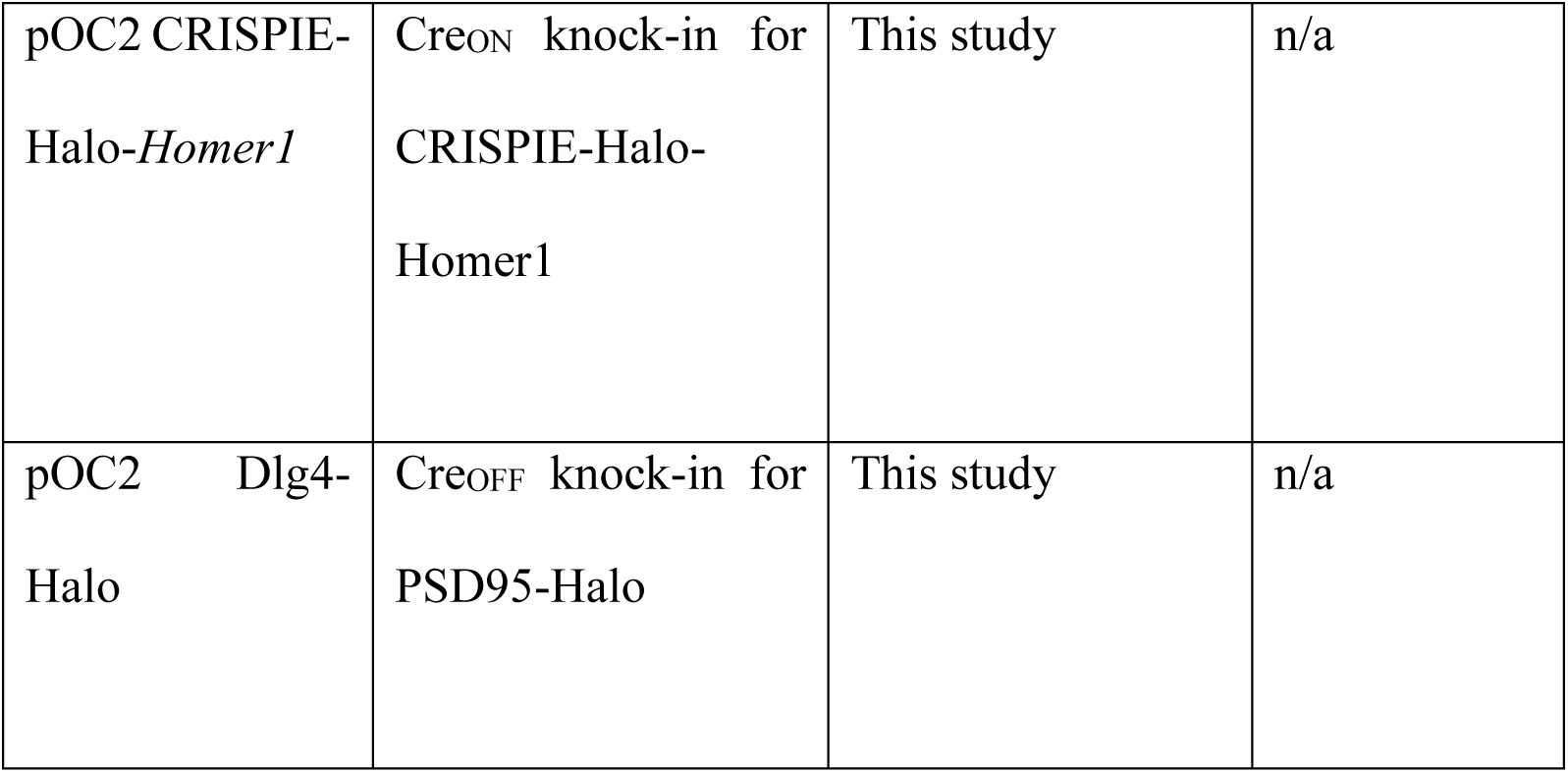
Vectors.

The Cre_ON_ knock-in vector (pOC1) is based on pORANGE LOX from (Willems *et al*., 2020, Addgene #139651), where CAG HA-SpCas9 was removed with XmaJI and NotI and replaced with primers AJ19164 and AJ19165 using primer ligation.

To clone the Cre_OFF_ knock-in vector (pOC2), lox-U6-lox was created by PCR from pOC1 with primers AJ19169 and AJ20047, digested with PscI and Bsp1407I, and ligated into pOC1 that was digested with PscI and Bsp1407I as well.

The Cre_ON_ Flp_OFF_ knock-in vector (pOC3) was based on pOC1, where one Frt site was cloned into the gRNA scaffold (Chylinski *et al*., 2019) and one before the U6 promoter. PCR fragments were obtained with primers AJ20124 – AJ20126, AJ20127 – AJ20125 and AJ20120 - AJ20121 using pOC1 as template. pOC1 was digested with NheI and PscI, and all fragments were ligated using HiFi assembly (New England Biolabs, Ipswich, United Kingdom).

The Flp_ON_ knock-in vector (pOC4) was designed analogous to Chylinski *et al*., 2019. PCR fragments were created using primers AJ20120 - AJ20121 with pOC1 as template, and primers AJ20122 – AJ20116 and AJ20117 – AJ20121 with pORANGE empty (Addgene #131471, Willems *et al*., 2020) as template. pORANGE was digested with PscI and NheI, and all fragments were ligated with HiFi assembly.

The Flp_OFF_ knock-in vector (pOC5) was designed similar as in Chylinski *et al*., 2019. PCR fragments were obtained with primers AJ21030 - AJ21031 with pOC3 as template, and AJ20122 and AJ21032 with pOC4 as template. pOC3 was digested with PscI and HindIII and fragments were ligated with HiFi assembly.

To obtain pORANGE β3-tubulin-2xGFP, the donor DNA from pORANGE β3-tubulin-GFP (Willems *et al*., 2020) was isolated using XmaJI and XbaI, and ligated in the XbaI site of pORANGE β3-tubulin-GFP. Subsequently, to create pORANGE β3-tubulin with 3xGFP or 4x-GFP, the pORANGE β3-tubulin-2xGFP double donor DNA was isolated with XmaJI and XbaI, and ligated in the XbaI site of pORANGE β3-tubulin-GFP or pORANGE β3-tubulin-2xGFP, respectively. Due to the repeated sequences, the orientation of the inserts could not be confirmed, but this should not affect performance of the knock-in.

pFUGW-NLS-Cre was created using a PCR reaction for NLS-Cre with primers AJ20128 - AJ20129 using pFUGW-GFP-NLS-Cre (a gift from Pascal Kaeser, Harvard Medical School Boston MA; Kaeser *et al*., 2011) as template. The PCR product was digested with NheI and XbaI, and ligated in NheI and XbaI sites of pFUGW.

pFUGW-NLS-Flp^O^ was created using a PCR reaction for NLS-Flp^O^ with primers AJ20130 - AJ20131 using pAAV hSynapsin Flp^O^ (Addgene #60663), a gift from Massimo Scanzianis (Xue, Atallah and Scanziani, 2014), as template. The PCR product was digested with NheI and XbaI, and ligated in NheI and XbaI sites of pFUGW.

pCAG ER^T2^-Cre-ER^T2^ lox was created by digesting pCAG ER^T2^-Cre-ER^T2^ (Matsuda and Cepko, 2007) with EcoRI and NotI. Both resulting DNA fragments were mixed with primers AJ21053 and AJ21054, and ligated with HiFi assembly.

All knock-ins were based on ORANGE (Suzuki *et al*., 2016; Willems *et al*., 2020) and cloned in pOCx backbones as described in Figure. S3B. gRNA target sequences can be found in Table 2. Fluorophores and epitope tags in the donor DNAs were exchanged using universal BmtI and AfeI restriction sites that are present in linker sequences surrounding the fluorophore. smFPs were obtained from Addgene #59759 (HA) and #59758 (V5), a gift from Loren Looger (Viswanathan *et al*., 2015).

**Table 2.**
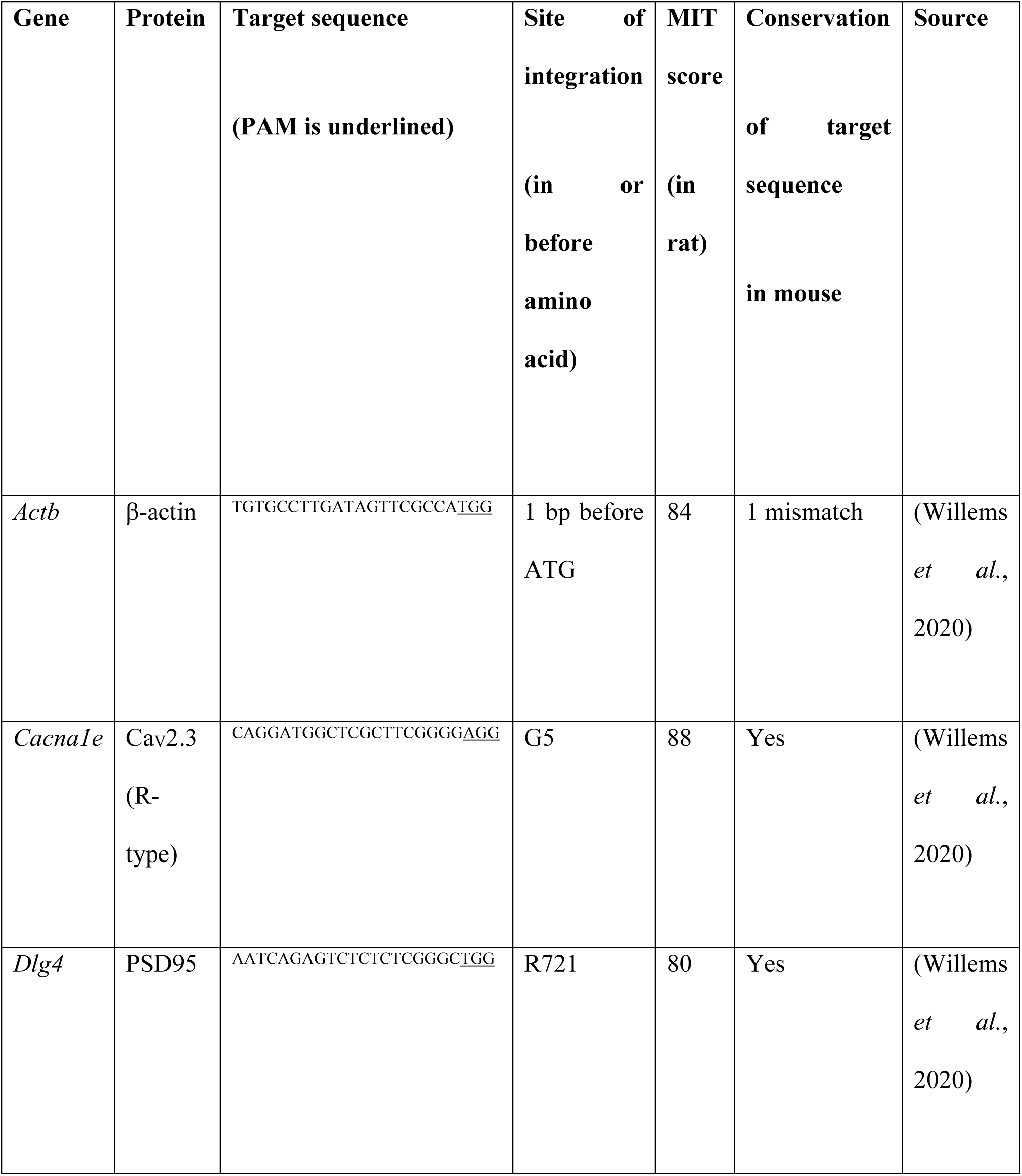
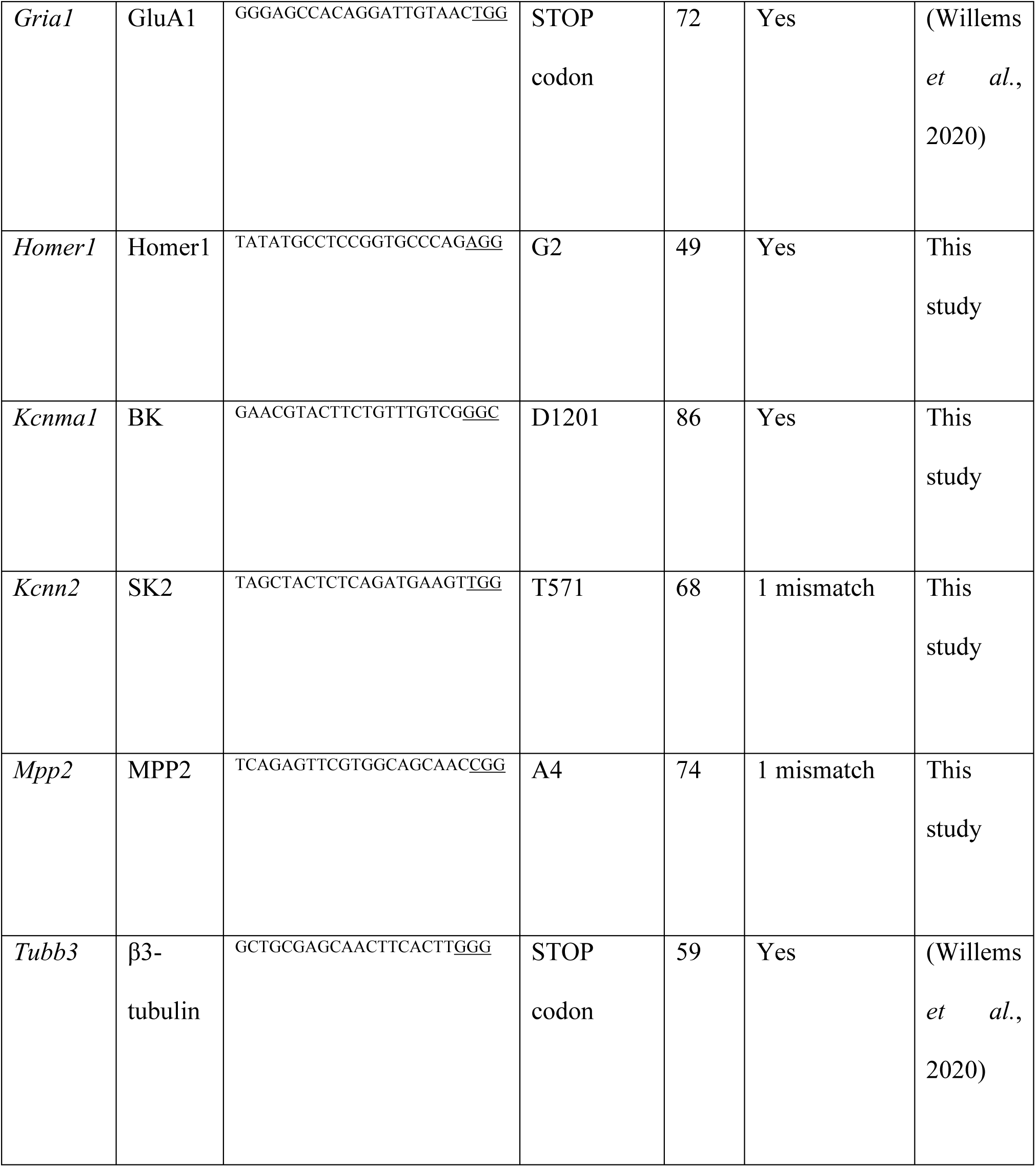
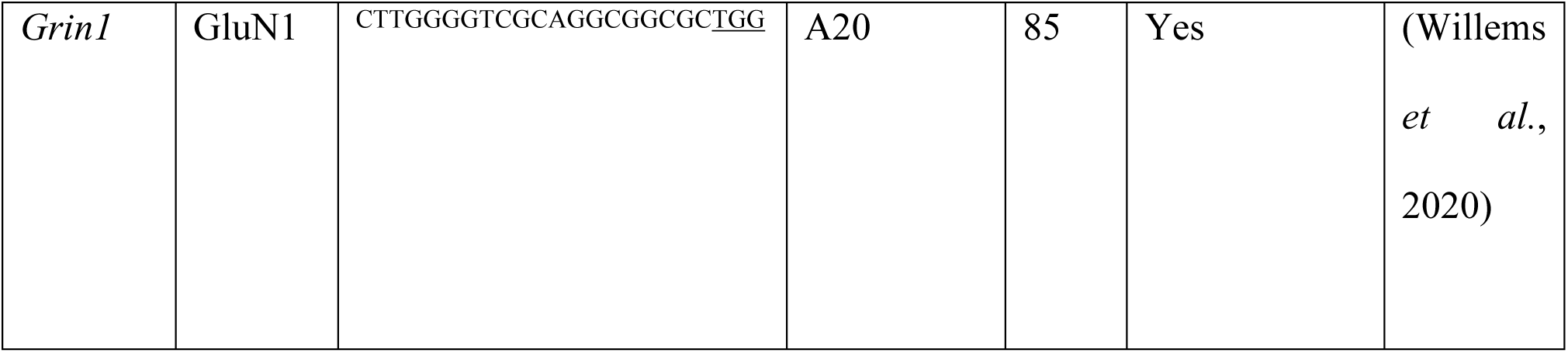
CRISPR knock-ins.

**Table 3.**
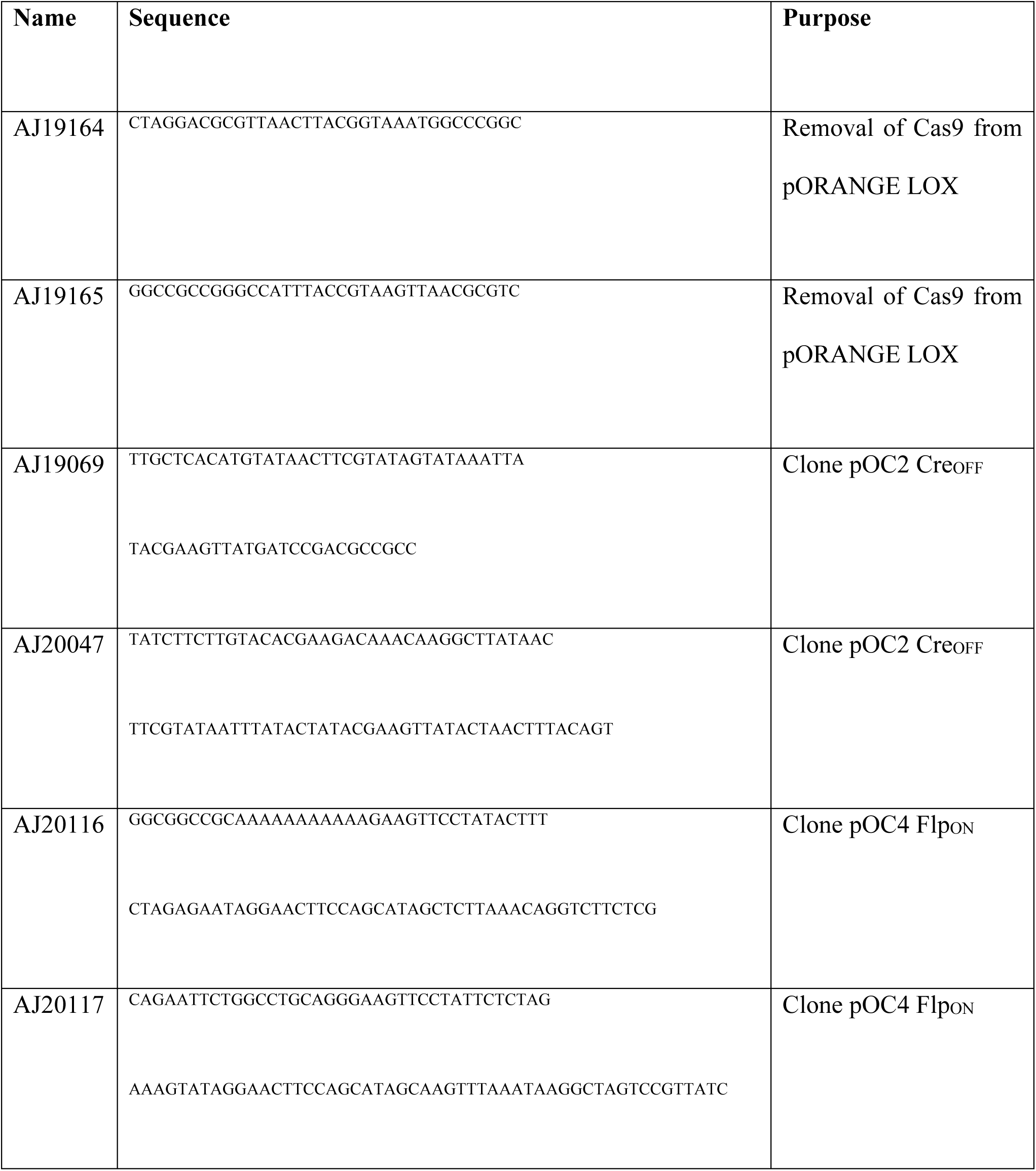
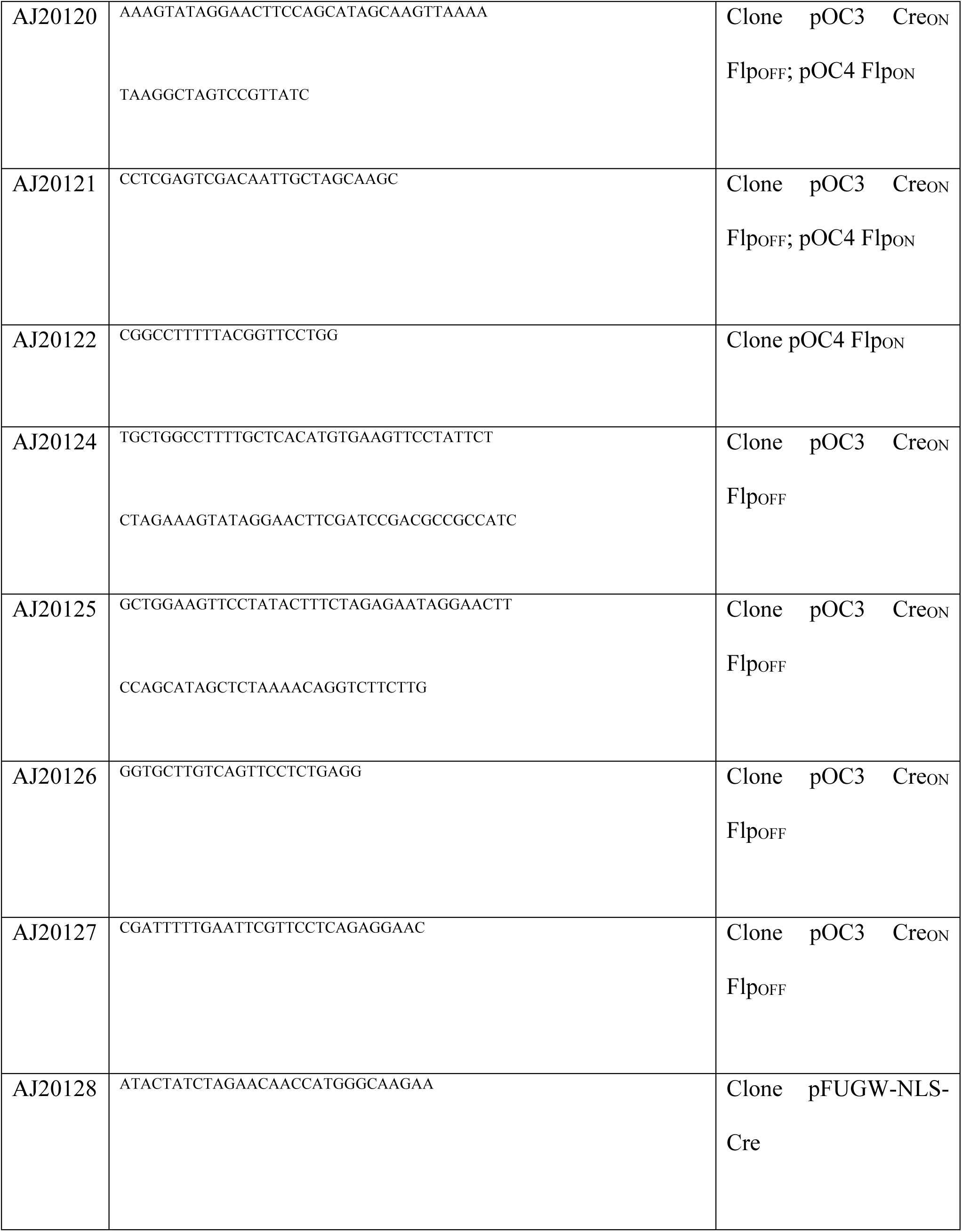
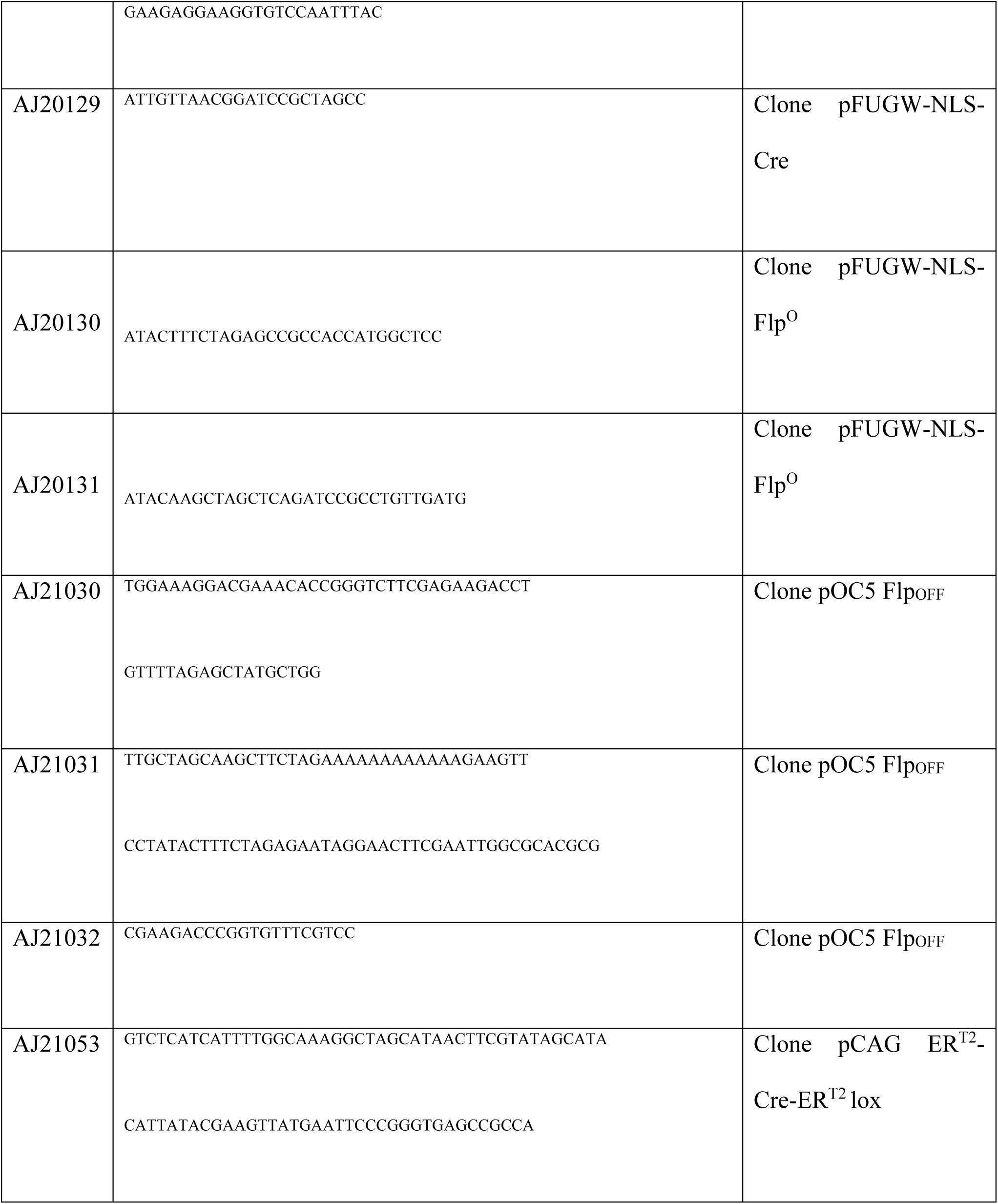
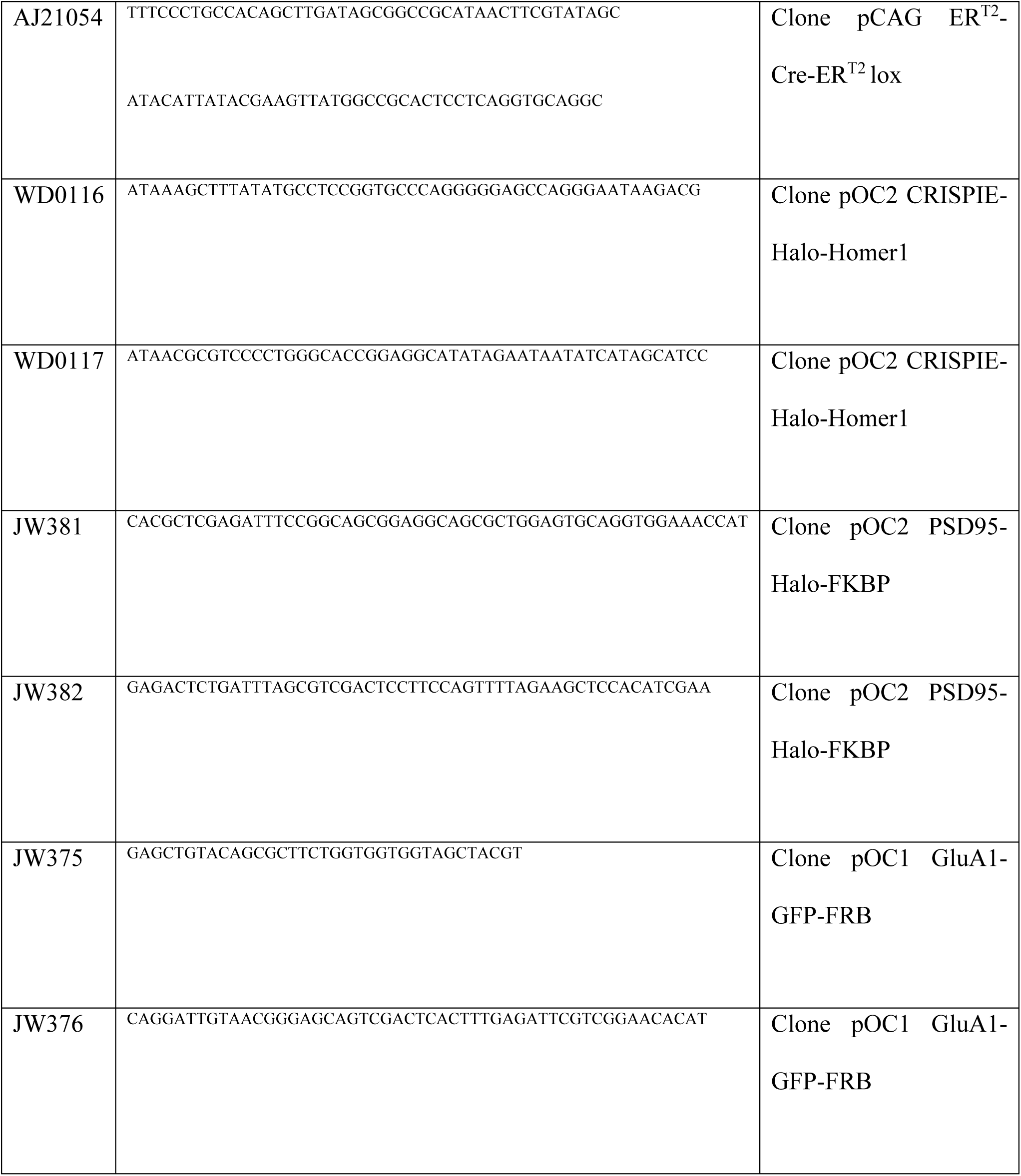
Primers.

For Rapalog-inducible assays, FKBP (primer JW381 ad JW382) and FRB (primer JW375 and JW376) were amplified using PCR (both a gift from Lukas Kapitein; Kapitein, Schlager, *et al*., 2010). These fragments were cloned into the AfeI site of pOC2 Dlg4-Halo and pOC1 Gria1-GFP, respectively, using HiFi assembly.

The design of pOC2 CRISPIE Halo-Homer1 was adapted from the strategy of Zhong *et al*., 2021 and described in Figure S4. The donor DNA was ordered as a gBlock from Addgene and PCR amplified using WD0116 and WD0117. Donor consists of Halo flanked by the splicing acceptor and donor of exon 8 of *Septin 3* (Fig. S4). Splicing acceptor and donor include 100 bp intronic and 10 bp exonic sequences. Target sequences were selected using CRISPOR and the BROAD institute sgRNA designer (Doench *et al*., 2016; Concordet and Haeussler, 2018).

### Antibodies and reagents

The following primary antibodies were used in this study: mouse anti-GFP 1:2000 dilution (Thermo Fisher, RRID: AB_221568), rabbit anti-GFP 1:2000 (MBL, RRID: AB_591819) rabbit anti-Halo 1:1000 (Promega, RRID: AB_713650), rat anti-HA 1:500 (Sigma-Aldrich, RRID: AB_390919) mouse anti V5 1:1000 (Invitrogen, RRID: AB_2556564). Alexa488-, Alexa568-, and Alexa647-conjugated secondary antibodies were used at 1:500 dilution, obtained from Life Technologies. CF568-conjugated secondary antibodies were used at 1:600 dilution, obtained from Sigma-Aldrich. Molecular biology reagents were obtained from Thermo Fisher. GFP minicircle for β3-tubulin were custom made at System Biosciences (Palo Alto, CA, USA), produced from pTubb3 MC (Addgene #87112, Suzuki *et al*., 2016). 4OH-tamoxifen was acquired from Sigma-Aldrich (cat. no H7904), and kept at 20 mM in ethanol in single use aliquots at -20°C. For Halo labeling, Halo Ligands JF549 (Promega, GA1110) and JF646 (Promega, GA1121) were used. Both dyes were dissolved using DMSO to 0.2 mM upon arrival. Working concentration was 0.2 µM (1:1000). For FKBP-FRB dimerization, we used Rapalog (TaKaRa, Shiga, Japan, #635057).

### Dissociated neuronal cultures

Dissociated hippocampal cultures were prepared from embryonic day 18 (E18) rat brains of both genders (Kapitein, Yau and Hoogenraad, 2010). Dissociated neurons were plated on Ø18mm coverslips coated with poly-L-lysine (37.5 µg/ml, Sigma-Aldrich) and laminin (1.25 µg/ml, Roche Diagnostics, Rotkreuz, Switzerland) at a density of 100,000 neurons per well, in Neurobasal medium (NB), supplemented with 1% penicillin and streptomycin (pen/strep), 2% B27, and 0.5 mM L-glutamine (all from Gibco;( NB-complete medium) at 37**°**C in 5% CO_2._ From DIV 1 onwards, medium was refreshed every 7 days by replacing half of the medium with Brainphys neuronal medium supplemented with 2% NeuroCult SM1 neuronal supplement (Stem cell technologies, Köln, Germany) and 1% pen/strep (BP-complete medium).

### Transfection of dissociated hippocampal neurons

Neurons were transfected at DIV 1-3 using Lipofectamine 2000 (Invitrogen). For one Ø18-mm coverslip seeded with 100,000 neurons, up to 2 μg DNA was used, which typically results in a few hundred to one thousand transfected cells per coverslip. DNA concentrations were determined using Nanodrop. For CAKE double knock-ins, the mixture contained 3.9 – 197 fmol (10-500 ng) Cre_OFF_ knock-in, 3.6 – 178 fmol (10-500 ng) Cre_ON_ knock-in, and 171 (500 ng) pCAG FLAG-SpCas9 expression. Experiments with inducible Cre used 2-20 fmol (10-100 ng) pCAG ERT2-Cre-ERT2 (Addgene #3777, Matsuda and Cepko, 2007) per coverslip. DNA was mixed with 3.3 μL Lipofectamine in 200 μL NB medium and incubated for 30 minutes at room temperature (RT). Next, 500 μL conditioned medium was transferred to a new culture plate and replaced by 300 μL NB supplemented with 0.5 mM L-glutamine. The DNA/Lipofectamine mix was added to the culture and incubated at 37°C and 5% CO2. After 90–120 minutes, coverslips were transferred to the new culture plate with conditioned medium and 500 μL new BP-complete and kept at 37°C and 5% CO2 for between 1–20 days, depending on the experiment.

### Immunocytochemistry

Hippocampal neurons were fixed at DIV 14-23 in 80 mM PIPES, 1 mM EGTA, 2 mM MgCl pH 6.8 and 4% paraformaldehyde (Electron Microscopy Sciences, Hatfield, PA, USA), for 5-10 minutes at 37°C and washed three times in PBS containing 0.1 M glycine (PBS/Gly). Neurons were blocked and permeabilized in blocking buffer (10% [v/v] normal goat serum [NGS] (Abcam, Cambridge, United Kingdom) in PBS/Gly with 0.1% [v/v] Triton X100) for 1 hour at 37°C. Next, coverslips were incubated with primary antibodies diluted in incubation buffer (5% [v/v] NGS in PBS/Gly with 0.1% [v/v] Triton X100) for 2 hrs at RT or overnight at 4°C, depending on the antibodies used. Coverslips were washed three times with PBS/Gly and incubated with secondary antibodies diluted 1:500 in incubation buffer for 1 hour at RT. Coverslips were washed three times in PBS/Gly, dipped in milliQ water (MQ), and mounted in Mowiol mounting medium (Sigma-Aldrich).

### Lentivirus production

For lentivirus production, HEK293T cells were maintained at a high growth rate in DMEM (Lonza, Basel, Switzerland) supplemented with 10% fetal calf serum (Corning) and 1% pen/strep. 1 day after plating on 10 cm dishes, cells at ∼70% confluency were transfected using Polyethylenimine (PEI, Polysciences. Warrington, PA, USA) with second-generation lentiviral packaging vectors (psPAX2 and 2MD2.G) and pFUGW-NLS-Flp^O^, pFUGW-NLS-Cre or pFUGW-GFP-NLS-Cre at a 1:1:1 molar ratio. At 6 hours after transfection, cells were washed once with PBS, and medium was replaced with 10 mL DMEM containing 1% pen/strep. At 48 hours after transfection, the supernatant was harvested, centrifuged 5 min at 700*g* to remove cell debris, and stored in aliquots at -80°C until use. Cultured hippocampal neurons were infected at DIV 3–9 with 20 μL virus added per well, unless indicated otherwise.

### Quantification of knock-in efficacy

To determine the efficacy of knock-ins, samples were fixed at DIV 14, and stained with primary and secondary antibodies as described above. Coverslips were examined with epifluorescence on a Nikon Eclipse 80i upright microscope Plan Fluor 20x air (N.A. 0.75) or Plan Fluor 40x oil (N.A. 1.30) objective, CoolLED pE-300^white^ illumination and Chroma ET-GFP/mCherry (59022) filter set. Coverslips were scanned top to bottom and fluorescent cells were scored manually based on staining pattern in one of four categories. For instance, for Cre_OFF_ β3-tubulin GFP / Cre_ON_ Halo-β-actin, the categories were 1) GFP correct (GFP staining pattern corresponds with β3-tubulin expression); 2) Halo correct (Halo staining pattern corresponds with β-actin pattern); 3) double correct (Both GFP and Halo have the correct staining pattern in the same cell); 4) incorrect (GFP staining pattern corresponds with β-actin and/or Halo corresponds with β3-tubulin). Results were obtained from 3 independent cultures with 1 coverslip per culture, unless stated otherwise.

### Confocal microscopy

Confocal images were acquired with a Zeiss LSM 700, using a EC Plan-Neofluar 40x oil (N.A.1.30) or Plan-Apochromat 63x oil (N.A. 1.40) objective and 488 nm, 555 nm and 633 nm laser excitation lines. Images were acquired as z-stacks containing planes at 0.5 µm interval, with 0.1 µm pixel size and 2x pixel averaging. All images are displayed as maximum intensity projections. Images in Figure S2A-B were acquired as tile scans, and stitched using Zeiss Zen Black 2.3 SP1 software.

### Spinning disc microscopy with FRAP

Neurons were transfected at DIV 2 as described above with pOC2 Dlg4-Halo-FKBP (33 fmol), pOC1 Gria1-GFP-FRB (33 fmol), pCAG ER^T2^-Cre-ER^T2^ (2 fmol) and pCAG FLAG-Cas9 (90 fmol). At DIV 7, 4OH-tamoxifen (100 nM) was added to the neurons. At DIV 21, just before imaging, neurons were live-labeled with Halo-ligand JF549 (diluted 1:1000 in conditioned medium) for 15 minutes and washed in conditioned medium for 10 minutes before mounting.

Imaging was performed on a spinning disk confocal system (CSU-X1-A1; Yokogawa, Musashino, Japan) mounted on a Nikon Eclipse Ti microscope (Nikon, Tokyo, Japan) with Plan Apo VC 100x 1.40 NA oil objective (Nikon) with excitation from Cobolt Calyspso (491 nm) And Cobolt Jive (561 nm), and emission filters (Chroma, ,Bellows Falls, VT, USA). The microscope was equipped with a motorized XYZ stage (ASI; MS-2000), Perfect Focus System (Nikon), Prime BSI sCMOS camera (Photometrics, Tucson, AZ, USA), and was controlled by MetaMorph software (Molecular Devices, San Jose, CA, USA). Neurons were mounted in a Ludin-chamber (Life Imaging Services) with 450 µL extracellular buffer (in mM: 140 NaCl, 3 KCl, 10 HEPES, 2.7 CaCl_2_, 2 MgCl_2_, and 10 D-glucose. pH 7.35) and maintained in a closed incubation chamber (Tokai hit: INUBG2E-ZILCS) at 37 °C.

Double knock-in neurons were imaged for 15 minutes, acquiring a Z-stack of 3 planes (0.5 µm interval) every 5 minutes. Hereafter, pre-selected ROIs (1.3 µm in diameter) on spines were bleached using the ILas2 system (Roper Scientific, Sarasota, FL, USA). After bleaching, the neurons were imaged every 5 minutes for a total of 30 minutes. Following the first acquisition, cells were incubated in 1 µM rapalog for 20 minutes. Next, a different part of the same neuron was selected for a second (after rapalog) round of FRAP imaging. As a control, Cre_ON_ GluA1-GFP-FRB positive neurons were imaged following the same protocol, but without the addition of rapalog.

Acquisitions were corrected for drift and a maximum projection of the Z-stack was used for analysis. For each ROI and timepoint, mean intensities were measured and corrected for background and bleaching. Mean intensities were normalized to 1 using the averaged intensities of the frames before bleaching, and normalized to 0 based on the intensity from the first frame after bleaching. The mobile fraction of GluA1-GFP-FRB was calculated by averaging the normalized intensity of the last 4 frames for each ROI. Analysis was performed using FIJI and Excel.

Synapse enrichment before and after rapalog was calculated as the ratio between synapse and dendritic shaft intensity using 10 ROIs (250 nm diameter) each. For this analysis, we used the images from timepoint -15 minutes.

### Dual-color SMLM

Neuron were transfected at DIV 1 as described above with pOC2 PSD95-GFP (7.9 fmol), pOC1 GluA1-HA (44 fmol), pCAG ERT2-Cre-ERT2 (2 fmol) and pCAG FLAG-Cas9 (90 fmol). 100 nM 4OH-tamoxifen was added at DIV 7. At DIV 23, cultures were fixed and stained with primary and secondary antibodies as described above. PSD95-GFP and GluA1-HA were labeled with Alexa647and CF568-conjugated secondary antibodies, respectively.

SMLM was performed on the Nanoimager S from ONI (Oxford Nanoimaging Ltd., Oxford, United Kingdom), fitted with a 100x 1.4NA oil-immersion objective, four laser lines (405 nm, 561 nm and 640 nm), an XYZ closed-loop piezo stage and a sCMOS camera (ORCA Flash 4, Hamamatsu). Integrated filters are used to split far-red emission onto the right side of the camera and blue-green-red emission spectra on the left side, enabling simultaneous dual-color imaging. The imaging chamber was temperature-controlled at 30°C to prevent fluctuations in temperature during the time course of an experiment that might affect the alignment of the channels. Channel alignment was performed before each imaging session using 100-nm TetraSpeck beads (T-7279, Invitrogen) and the ONI software aiming for an alignment error of SD < 7 nm as measured from 2000 points total across a maximum of 20 fields of view. Imaging was performed in near-TIRF using a motorized mirror. During acquisition, neurons were kept in a STORM buffer (pH 8.0) containing 50 mM Tris, 10 mM NaCl, 10% w/v D-glucose, 5 mM MEA, 700 μg/ml glucose oxidase, and 40 μg/ml catalase. For each double knock-in neuron, 20,000 frames were acquired at 50 Hz. NimOS software from ONI was used for detection of single molecule localization events. Resulting localization tables were drift-corrected using Detection of Molecules (DoM) plugin v.1.2.1 for ImageJ (https://github.com/ekatrukha/DoM_Utrecht). dSTORM reconstruction was made using DoM with pixel size of 12 x 12 nm (Fig. 6A). Analysis was continued in MATLAB (2021a).

Localizations were filtered out if localization precision was > 30 nm for GluA1 and > 25 nm for PSD95, or photon count was < 300 or > 30,000 photons. Consecutive localizations in a radius of 60 nm were removed. If consecutive localizations persisted for more than 10 frames, the initial localization was also removed. ROIs outlining the synapse were defined based on the full width half maximum (FWHM) using a widefield image of PSD95-GFP taken before dSTORM acquisition. Synapses were only analyzed further if out if they contained > 800 localizations for PSD95 and > 400 localizations for GluA1, or if they were > 0.02 μm^2^ or < 0.3 μm^2^ in size. For each localization in a given synapse, the local density was calculated as the number of localizations within a radius of 5 * the mean nearest neighbor distance (MacGillavry *et al*., 2013). Localizations were deemed part of a nanodomain if its local density was > 40. Nanodomains were isolated using the MATLAB functions linkage() and cluster(). Subsequently, nanodomains were subclustered if they contained multiple local density peaks that were > 80% of the maximum local density, further than 80 nm apart and separated by a local minimum of < 30% of the maximum local density. The nanodomain boundary was constructed using the Voronoi diagrams circumventing the localizations. Nanodomains with < 5% of the localizations in a synapse or a diameter of < 30 nm were excluded. The center of the PSD was calculated using the function centroid() and based on the PSD95 cluster inside the synapse, identified using DBSCAN (Ester *et al*., 1996). Nanodomain distance between PSD95 and GluA1 was calculated for each nanodomain as the distance to its closest nanodomain in the other channel. Co-localization analysis of PSD95 and GluA1 was performed as described in Willems & MacGillavry (*in review)*. As a first step in determining the co-localization between two clusters, the local density is determined for each localization in both channels. The mean nearest neighbor distance (MNND) is determined within the PSD using the MATLAB function *knnsearch*. Next, for each localization, the local density (*LD*) is defined as the number of localizations within a radius defined by the effective resolution making use of the MATLAB function *rangesearch*. For each channel, the *LD* values are averaged together to obtain *LD_A_* and *LD_B_.* Effective resolution was calculated as (Gould, Verkhusha and Hess, 2009):

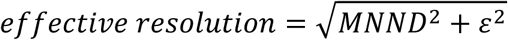

where *ε* is the localization error.

The co-localization index is determined as the number of localizations of channel B (*N*) within a radius (*d*) around each localization in channel A (*Ai*) normalized to the mean *LD* of the localizations in channel 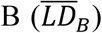, with *d* being the effective resolution of the localizations in channel B:

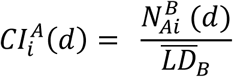

Thus, for channel B, the co-localization index values are calculated as:

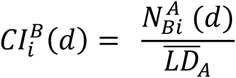

The co-localization indices calculated for each localization individually are used to plot a co-localization map and averaged to obtain a mean co-localization index per synapse for both channels (Fig. 6E).

### Data representation and statistics

All experiment were performed in at least 3 independent cultures. Data is shown as average values, error bars represent standard error of the mean (SEM). When comparing two experimental groups, an unpaired t-test was used, except for the FRAP analysis, which was analyzed with a paired t-test. If groups were not normally distributed, the non-parametric Mann-Whitney test was used. Cell counting experiments were analyzed with a 1-way or 2-way ANOVA, and for results with p < 0.05, a post hoc test with Tukey-Kramer correction was performed to test for differences between individual groups.

## Acknowledgements

This work was supported by the European Research Council (ERC-StG 716011 to HDM), NWO (OCENW.KLEIN.163 to HDM) and a NARSAD Young Investigator Grant from the Brain & Behavior Research Foundation (grant number 29452) to AdJ. We thank Manon Westra for contribution to single molecule localization scripts, and Martin Harterink for critically reading the manuscript.

## Author contributions

Conceptualization: AdJ

Methodology: AdJ

Software: WD, JW

Formal analysis: WD, JW,

AdJ Investigation: WD, JW, AdJ

Writing – original draft preparation: AdJ

Writing – Review & editing: WD, JW, HDM

Visualization: WD, JW, AdJ

Supervision: AdJ, HDM

Funding Acquisition: AdJ, HDM

## Supplemental Figures

**Figure S1.**
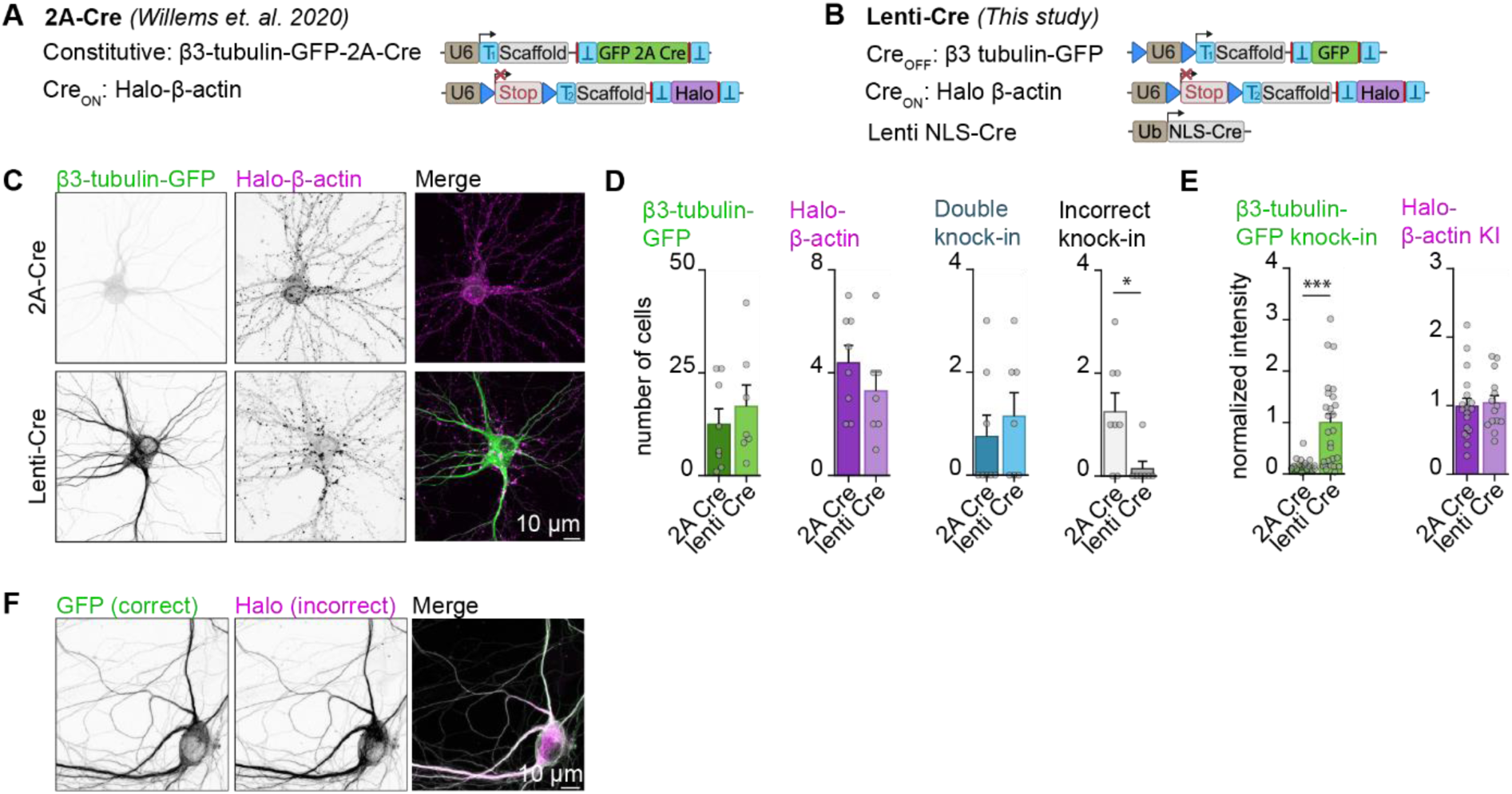
Improved CAKE mechanism reduces crosstalk between knock-ins. **A.** Overview of original CAKE knock-in mechanism introduced in (Willems *et al*., 2020), referred to as 2A-Cre throughout this Figure. The constitutively active knock-in labels β3-tubulin with GFP-2A-Cre. Cre expression from this allele then activates gRNA expression for the Cre_ON_ Halo-β-actin knock-in. **B.** Overview of improved CAKE constructs introduced in this study (see main text) which will be referred to as Lenti-Cre throughout this Figure. Lentivirus encoding for Cre is added at DIV 7, which switches off gRNA expression for Cre_OFF_, and switches on gRNA expression for Cre_ON_. **C.** Example confocal images of CAKE double knock-ins using the 2A-Cre or Lenti-Cre mechanism. **D.** Average number of single and double knock-in cells per coverslip for 2A-Cre and Lenti-Cre CAKE. 2A-Cre n = 8 coverslips, lenti-Cre n = 7 coverslips, N = 4 independent cultures. β3-tubulin-GFP *p* = 0.50, Halo-β-actin *p* = 0.29, double knock-in *p* = 0.53, incorrect knock-in * *p* = 0.019, unpaired *t*-test. **E.** Average fluorescence intensity in proximal dendrites of single knock-ins, normalized per culture. For β3-tubulin-GFP: 2A-Cre n = 21 cells, lenti-Cre n = 25 cells, N = 3 independent cultures. For Halo-β-actin: 2A-Cre n = 18 cells, lenti-Cre n = 14 cells, N = 3 independent cultures. β3-tubulin-GFP * *p* = 0.001, Halo-β-actin p = 0.29, unpaired *t*-test. **F.** Example confocal image of an incorrect double knock-in. Fluorescence signal of both GFP and Halo are consistent with β3-tubulin distribution, suggesting that the Halo donor was inserted in *Tubb3*.

**Figure S2.**
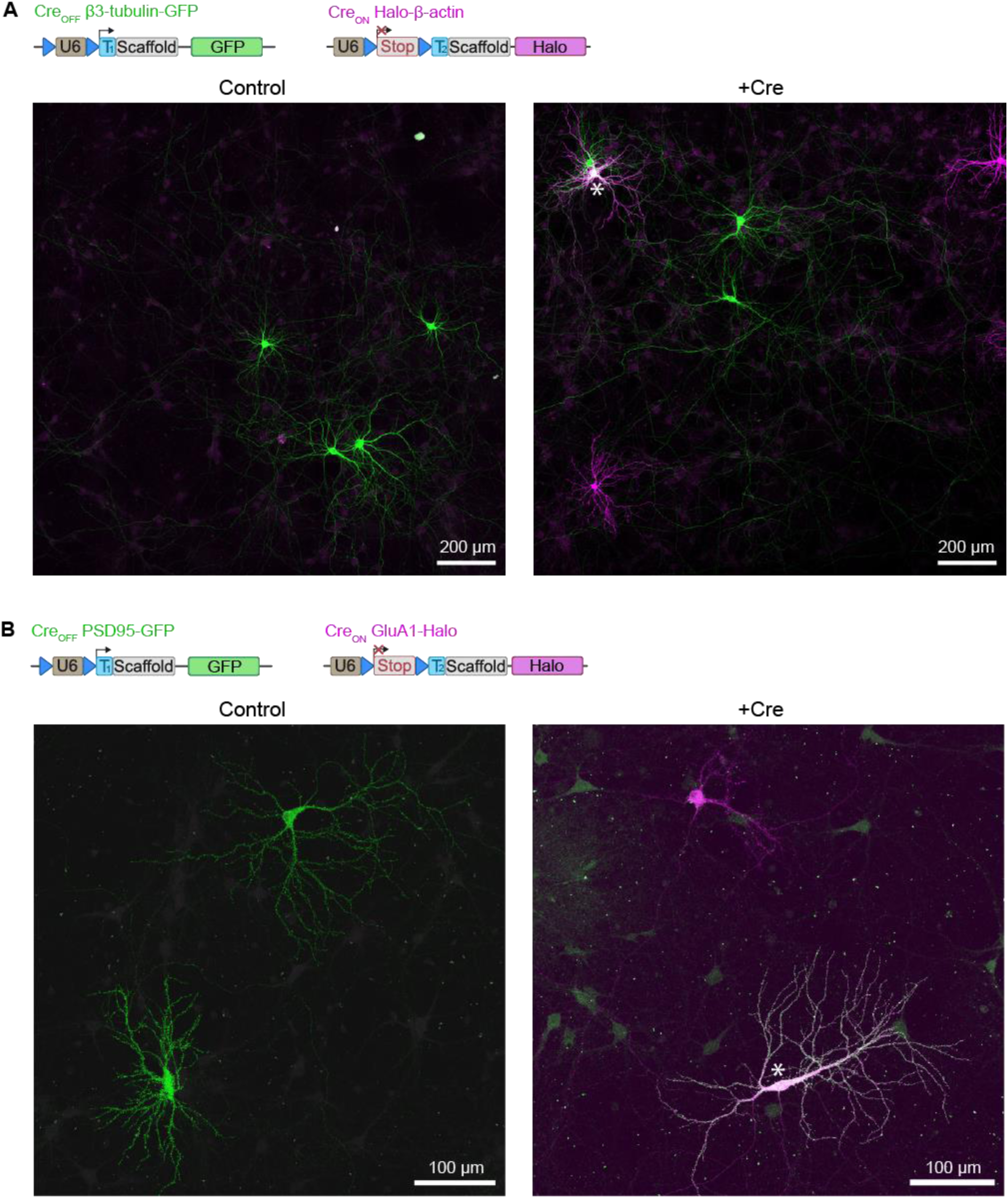
Low magnification images of CAKE knock-in cultures. **A.** Low magnification images of DIV 14 culture transfected with Cre_OFF_ β3-tubulin-GFP and Cre_ON_ Halo-β-actin knock-in vectors. Control is unstimulated, +Cre condition was infected with 20 µL lenti-Cre at DIV 7. Notice that CAKE leads to a mosaic of single and double knock-ins (indicated with *). Fluorescent neurons are surrounded by untransfected cells. **B.** As in A, for Cre_OFF_ PSD95-GFP and Cre_ON_ GluA1-Halo.

**Figure S3.**
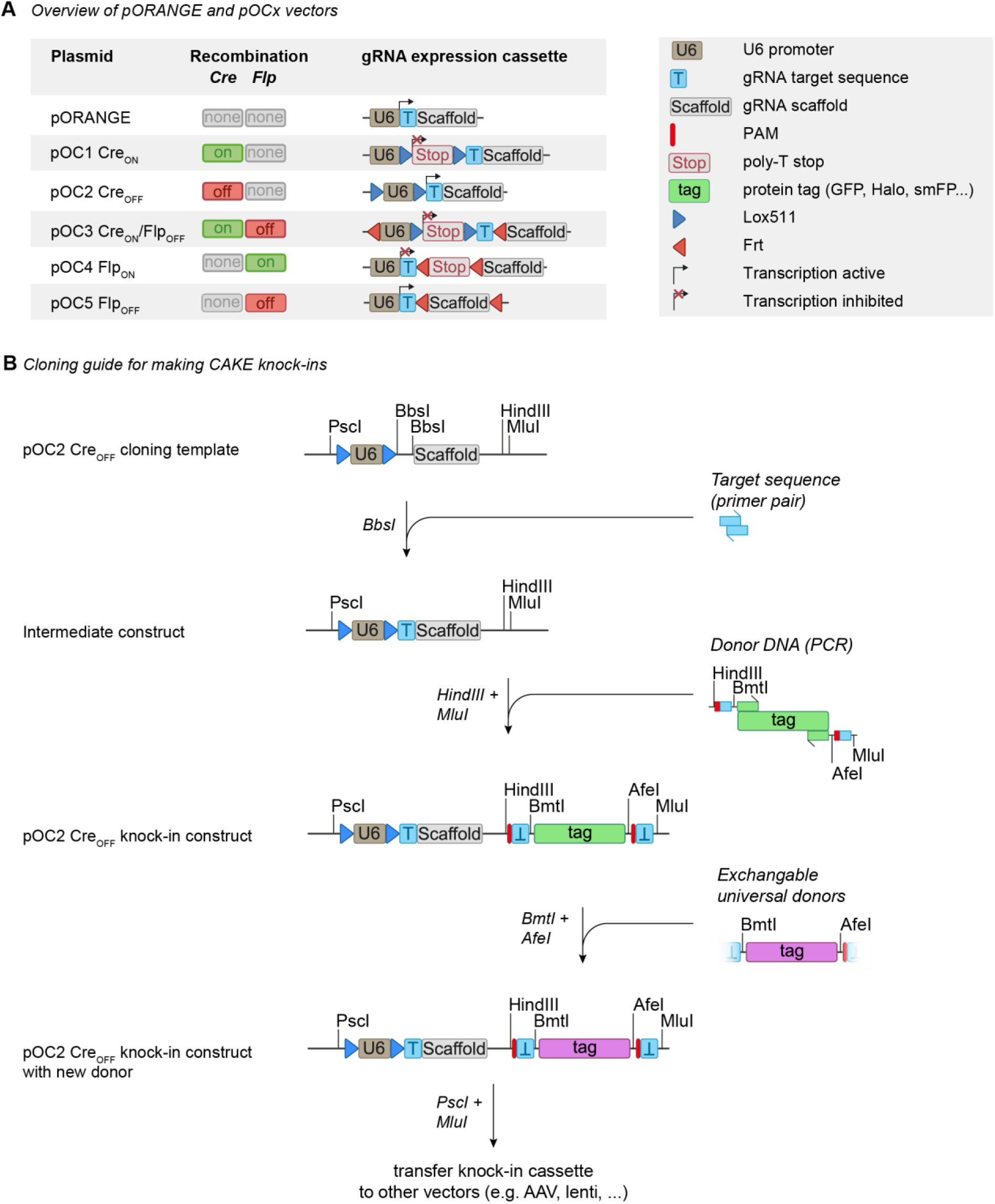
Overview and cloning guide for CAKE template vector library. **A.** Overview of pORANGE (Willems *et al*., 2020) and the CAKE template vectors introduced in this study. All vectors contain a multiple cloning site for addition of the donor DNA. **B.** Cloning guide to create CAKE knock-in constructs. The example shows a step-by-step protocol for pOC2 Cre_OFF_ knock-in vectors, but identical cloning steps apply to all pOC vectors shown in A. #1 shows the empty pOC2 template vector. After digestion with BbsI, a primer pair encoding for the gRNA target sequence is added to obtain the intermediate construct (#2). The donor DNA, which always contains a protospacer adjacent motif (PAM) and target sequence (Suzuki *et al*., 2016) is obtained with standard PCR techniques. Figure S1 of Willems *et al*., (2020) contains a detailed description on design of the donor DNA. The donor is cloned using HindIII and MluI restriction sites to obtain the final pOC2 knock-in construct (#3). Most of the knock-in constructs used in this study contain BmtI and AfeI restriction sites around the fluorophore, for universal exchange of donors between knock-in vectors, without the need of PCR (shown in #4, optional). The entire knock-in cassette, containing U6-driven gRNA expression, Lox551 and/or Frt sites and the donor DNA, can be removed from pOC vectors using PscI and MluI restriction sites.

**Figure S4.**
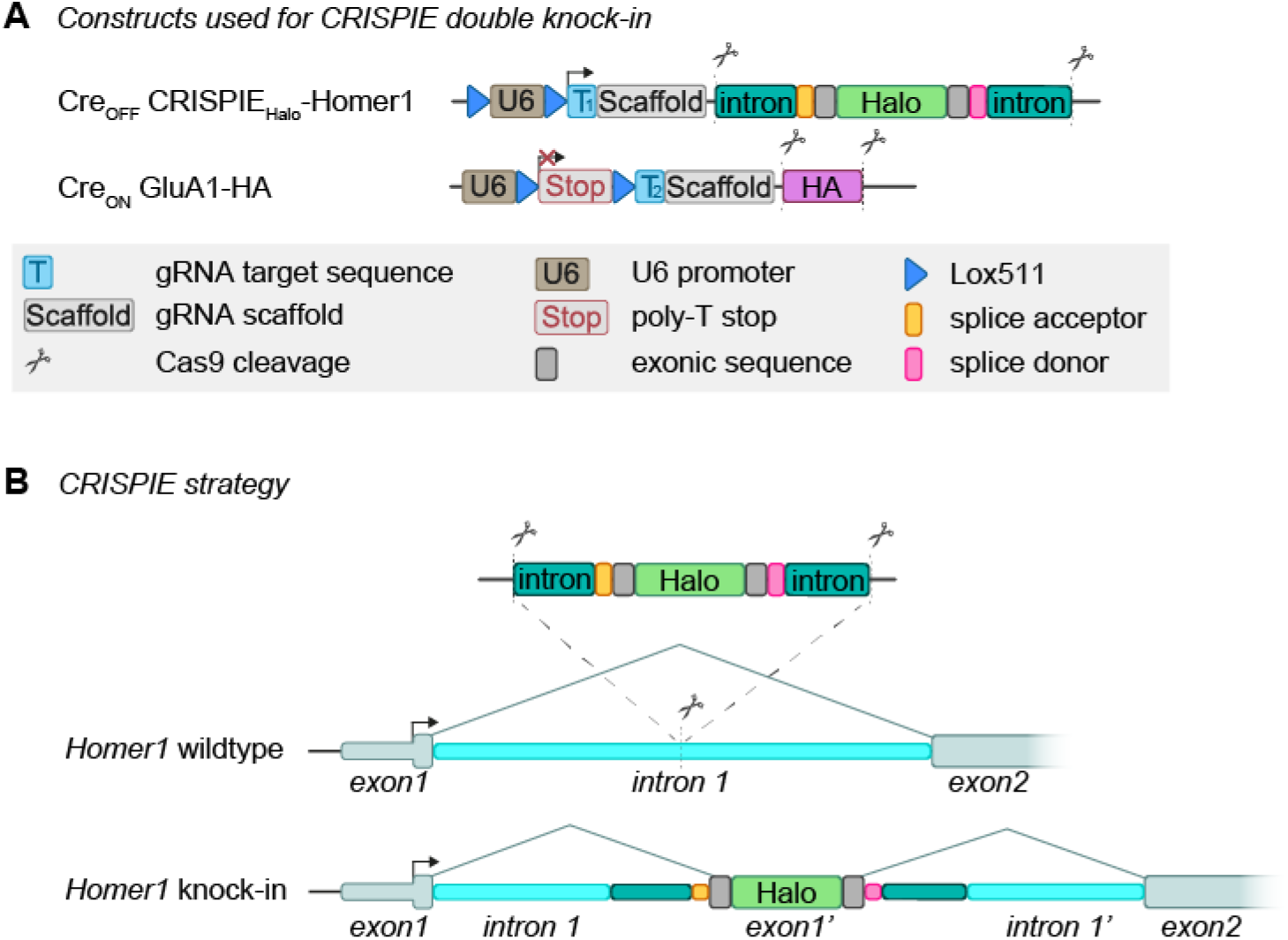
Overview of CRISPIE strategy. **A.** Overview of DNA constructs used for the CRISPIE double knock-in. **B.** Visual representation of the CRISPIE modality. Halo, flanked by the splicing acceptor and donor sites of exon 8 of *Septin3*, is inserted into the first intron of *Homer1* following Cas9 cleavage. Halo then functions as exon1’ and will be integrated in the mRNA.

**Figure S5.**
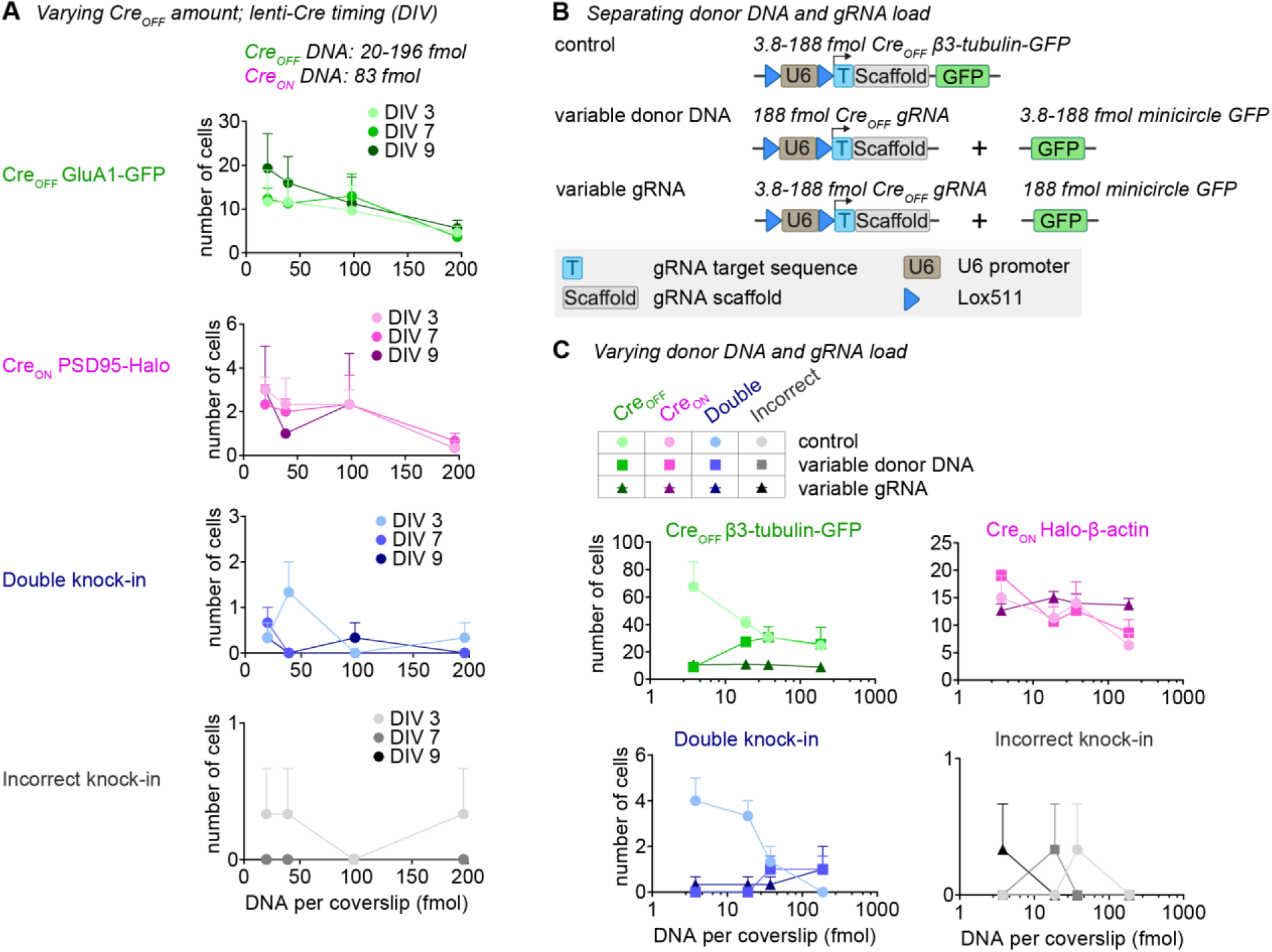
Donor DNA amount controls the number of knock-ins. **A.** Number of fluorescent cells per coverslip for each knock-in, as a function of Cre_OFF_ GluA1-GFP vector amount. 20 µL lenti-Cre was added at DIV 3, 7 or 9. n = 3 coverslips, N = 3 independent cultures. For incorrect knock-ins, only double-incorrect cells were scored. **B.** Overview of conditions and constructs. Cre_OFF_ β3-tubulin-GFP is the same vector as used in the main text (for instance in Fig. 1). The Cre_OFF_ gRNA is the same vector, but without the GFP donor DNA. Minicircle donor encodes for the GFP donor only, with a single protospacer adjacent motif and target sequence (Suzuki *et al*., 2016). **C.** Number of fluorescent cells per coverslip for each knock-in. ER^T2^-Cre-ER^T2^ was used for this experiment, and 100 nM 4OH-tamoxifen was added at DIV 7. n = 3 coverslips, N = 3 independent cultures.

**Figure S6.**
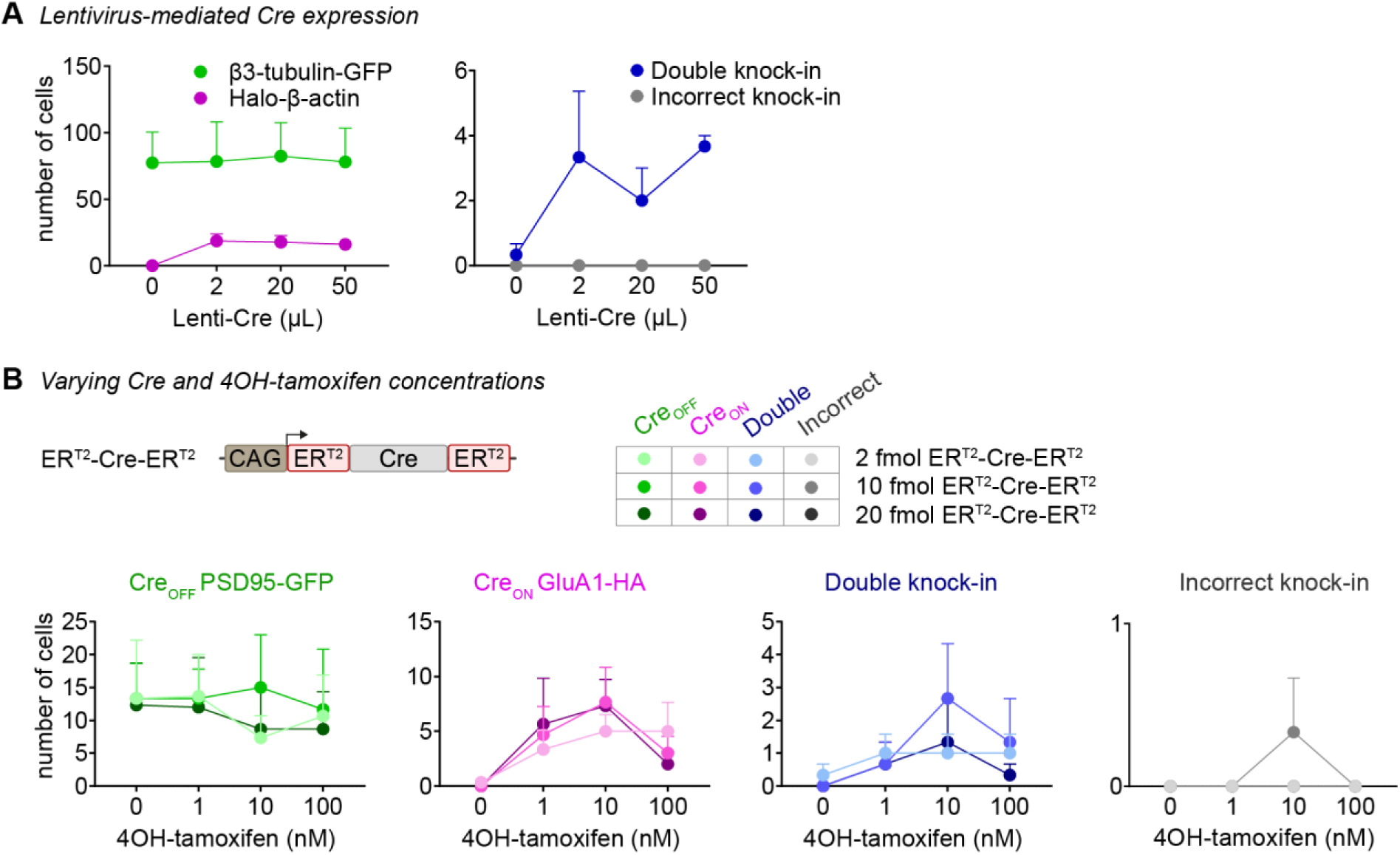
Optimization of lenti-Cre and ER^T2^-Cre-ER^T2^. **A.** Number of fluorescent cells per coverslip as a function of the amount of lenti-Cre used for infection. Experiments were perfomed with Cre_OFF_ β3-tubulin-GFP and Cre_ON_ Halo-β-actin. Lenti-Cre was added at DIV 7. n = 2 coverslips N = 2 independent cultures. **B.** Number of fluorescent cells per coverslip as a function of 4OH-tamoxifen concentration and amount of pCAG ER_T2_-Cre-ER^T2^ vector at transfection. At 2 fmol (10 ng) pCAG ER^T2^-Cre-ER^T2^, 4OH-tamoxifen results in a dose dependent increase in the number of Cre_ON_ knock-ins. Higher vector load (10-20 fmol) does not lead to more knock-in cells, but increases ER^T2^-Cre-ER^T2^ cytotoxicity at 100 nM 4OH-tamoxifen. n = 3 coverslips, N = 3 independent cultures.

**Figure S7.**
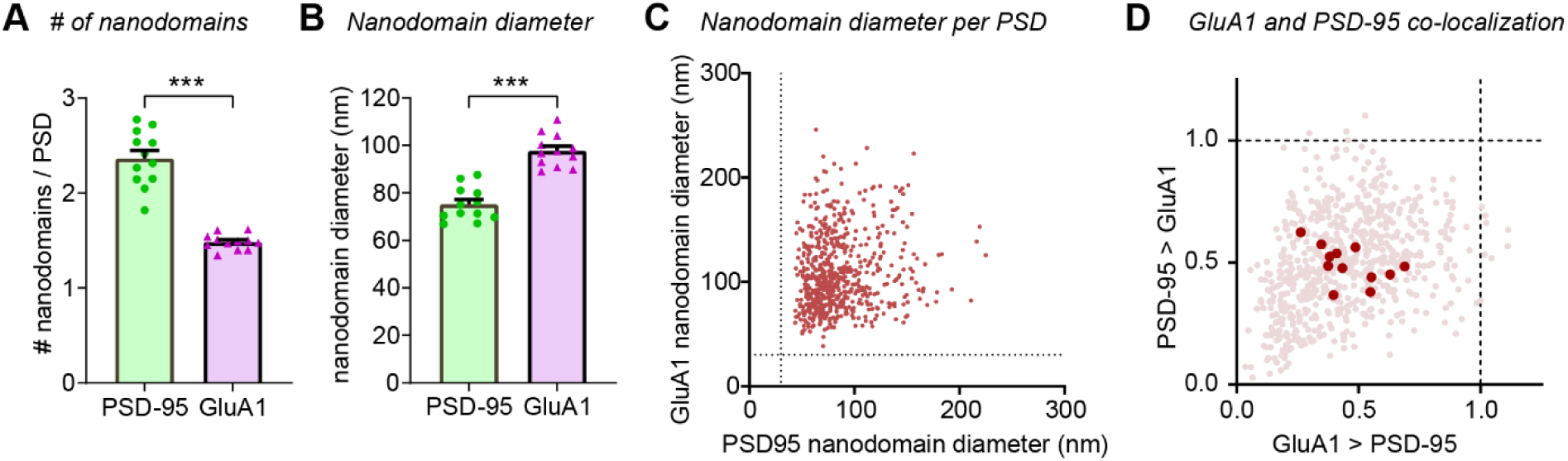
PSD95 and GluA1 have distinct nanoscale organizations. **A.** Number of nanodomains of PSD95 and GluA1 per PSD. *** *p* = 0.0002, unpaired *t*-test. **B.** Nanodomain diameter for PSD95 and GluA1. *** *p* = 6.4 * 10^-8^, unpaired *t*-test. **C.** Nanodomain diameter for PSD95 and GluA1 per PSD. Dotted line represents minimal threshold of nanodomain diameter. **D.** Co-localization index for both PSD95 and GluA1, plotted per synapse (transparent red dots) and neuron (red dots).

**Figure S8.**
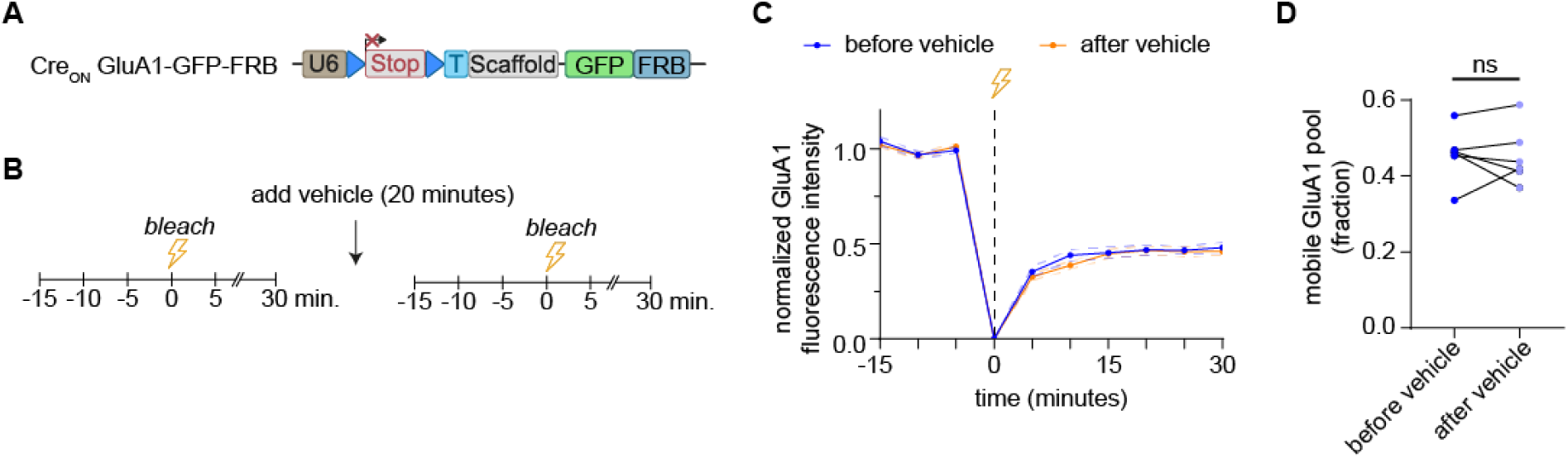
Live-cell anchoring of AMPA receptors control. **A.** Overview of DNA construct used. **B.** Imaging protocol. Neurons are live-cell imaged using spinning disk microscopy every 5 minutes. FRAP is performed twice, before and after vehicle incubation for 20 minutes. **C.** FRAP curves of spines bleached before and after vehicle incubation. Data is normalized to the average intensity before bleaching. 34 spines before vehicle and 42 spines after vehicle were bleached. n = 6 cells, N = 5 independent cultures. **D.** Average recovery of fluorescence per neuron, averaged over the last 4 frames, reflecting the mobile pool of receptors. No difference was observed before and after vehicle. *p* = 0.86, paired *t*-test. N = 6 cells, N = 5 independent cultures.

